# Purine and carbohydrate availability drive *Enterococcus faecalis* fitness during wound infection

**DOI:** 10.1101/2022.10.17.512645

**Authors:** Casandra Ai Zhu Tan, Kelvin Kian Long Chong, Daryl Yu Xuan Yeong, Celine Hui Min Ng, Muhammad Hafiz Ismail, Vanessa Shi Yun Tay, Yusuf Ali, Kimberly A. Kline

## Abstract

*Enterococcus faecalis* is commonly isolated from a variety of wound types. Despite its prevalence, the pathogenic mechanisms of *E. faecalis* during wound infection are poorly understood. Using a mouse wound infection model, we performed *in vivo E. faecalis* transposon sequencing and RNA sequencing to identify fitness determinants that are crucial for replication and persistence of *E. faecalis* during wound infection. We found that *E. faecalis* purine biosynthesis genes are important for bacterial replication during the early stages of wound infection, a time when purine metabolites are rapidly consumed by *E. faecalis* within wounds. We also found that the *E. faecalis* MptABCD phosphotransferase system (PTS), involved in the import of galactose and mannose, is crucial for *E. faecalis* persistence within wounds of both healthy and diabetic mice, especially when carbohydrate availability changes throughout the course of infection. During *in vitro* growth with mannose as the sole carbohydrate source, shikimate and purine biosynthesis genes were downregulated in the OG1RF Δ*mptD* mutant compared to the isogenic wild-type strain, suggesting a link between mannose transport, shikimate, and purine biosynthesis. Together, our results suggest that dynamic and temporal microenvironment changes at the wound site affects pathogenic requirements and mechanisms of *E. faecalis* and raise the possibility of lowering exogenous purine availability and/or targeting galactose/mannose PTS to control wound infections.

**IMPORTANCE:** Although *E. faecalis* is a common wound pathogen, its pathogenic mechanisms during wound infection are unexplored. Here, combining a mouse wound infection model with *in vivo* transposon and RNA sequencing approaches, we identified the *E. faecalis* purine biosynthetic pathway and galactose/mannose MptABCD phosphotransferase system as essential for *E. faecalis* acute replication and persistence during wound infection, respectively. The essentiality of purine biosynthesis and the MptABCD PTS is driven by the rapid consumption of purine metabolites by *E. faecalis* during acute replication and changing carbohydrate availability during the course of wound infection. Overall, our findings reveal the importance of the wound microenvironment in *E. faecalis* wound pathogenesis and how these metabolic pathways can be targeted to better control wound infections.

## INTRODUCTION

Wound infections affect approximately 11 million people worldwide (Demidova-Rice et al., 2012) and roughly 20 billion USD is spent yearly on treatment (Sen et al., 2009). Wounds are broadly classified as acute or chronic. Although acute wounds heal in a predictable time course while chronic wounds are perturbed during the wound healing process(es), both are prone to colonization by a diversity of microorganisms such as enterococci (Bowler et al., 2001). *Enterococcus faecalis* is a common pathogen that can colonize different wound types ranging from surgical sites, chronic ulcers, and diabetic wounds (Dowd et al., 2008; Dworniczek et al., 2012; Fisher & Phillips, 2009; Giacometti et al., 2000; Shettigar et al., 2018). Moreover, *E. faecalis* is highly resilient to environmental stressors such as a broad pH range and high salt conditions (Arias & Murray, 2012; Flahaut et al., 1996) and form antibiotic tolerance-associated biofilm microcolonies on the wound bed (Chong et al., 2017; James et al., 2008); together rendering *E. faecalis* wound infection difficult to treat.

Despite many studies undertaken to investigate the pathogenic mechanisms of *E. faecalis* in other infection sites, little is known regarding the pathogenicity of *E. faecalis* during wound infection. We previously showed, using a mouse wound excisional infection model, that *E. faecalis* undergoes acute replication within wounds during the first 8 h post-infection (hpi), followed by a continuous tapering of bacterial colony-forming unit (CFU) until 3 days post-infection (dpi), after which the bacterial load was maintained at 10^5^ CFU until 7 dpi (Chong et al., 2017). In the same study, *E. faecalis* multiple peptide resistance factor (MprF), which confers protection against host-derived cationic antimicrobial peptides (CAMPs) (Bao et al., 2012; Kandaswamy et al., 2013; Peschel et al., 2001; Samant et al., 2009; Thedieck et al., 2006), was identified as a fitness determinant involved in *E. faecalis* persistence in wounds at 3 dpi (Chong et al., 2017). To date, besides MprF, no other fitness (or metabolic) determinants have been identified that contribute to *E. faecalis* replication (8 hpi) and persistence (3 dpi) during wound infection.

To address this knowledge gap, we used *in vivo* transposon and RNA sequencing to probe for additional fitness determinant(s) that contribute to replication and persistence during *E. faecalis* wound infection. We identified *de novo* purine biosynthesis genes to be indispensable for *E. faecalis* acute replication during wound infection. Liquid chromatography-mass spectrometry (LC-MS) analysis of mouse wound samples showed that exogenous purine metabolites in the wound microenvironment are rapidly consumed by *E. faecalis*, and thus likely insufficient to support *E. faecalis* rapid growth during wound infection. We also identified the *E. faecalis* MptABCD phosphotransferase system (PTS) to be crucial for persistence in wounds. We characterized the *E. faecalis* MptABCD PTS as a galactose and mannose transporter, and found that carbohydrate availability such as glucose, galactose, and mannose changes as wound infection progress. In addition to a role during wound infection pathogenesis, we also showed that both *E. faecalis de novo* purine biosynthesis and MptABCD PTS contribute to *E. faecalis* fitness during catheter-associated urinary tract infection (CAUTI). Altogether, our study suggests that changes in the wound microenvironment affects *E. faecalis* pathogenesis and raises the possibility of reducing purine availability in the wound microenvironment and/or targeting MptABCD PTS as future therapeutic targets to curb chronic wound infections.

## MATERIALS AND METHODS

### Bacterial strains and growth conditions

Bacterial strains used in this study are listed in **Supplementary Table 1**. Unless stated, all *E. faecalis* bacterial strains were grown in Brain Heart Infusion broth (BHI; Neogen, USA) at 37 °C in static conditions for 16 – 18 h. Cells were harvested by centrifugation at 5,000 rpm for 5 min and cell pellets were washed twice with 1 mL of 1X sterile phosphate buffered saline (PBS). The final pellet was resuspended in 5 mL of 1X sterile PBS prior to optical density (OD) measurement at 600 nm. Cell suspensions were then normalized to the required cell number for the various experimental assays. When applicable, BHI were supplemented with 25 µg/mL erythromycin (Sigma-Aldrich, USA) for maintenance of pTCV and pMSP3535 plasmids.

### Mouse wound excisional model

Bacterial cultures were normalized to 2 – 4 × 10^8^ CFU/mL in 1X sterile PBS. Mouse wound infections were performed similar to a previous study (Chong et al., 2017). Briefly, male C57BL/6 mice (7 – 8 weeks old, InVivos, Singapore) or male and female *db/db* (BKS.Cg-*Dock7^m^ +/+ Lepr^db^/*J) mice (7 – 8 weeks or 14 weeks old, The Jackson Laboratory, USA) were anesthetized by inhalation of 3% isoflurane and the dorsal hair trimmed. A depilatory cream (Nair cream, Church and Dwight Co, USA) was then applied, and fine hair was removed through shaving with a scalpel. The skin was subsequently disinfected with 70% ethanol and a wound was created using a 6 mm biopsy punch (Integra Miltex, USA). This was followed by inoculation with 10 μL of respective bacterial cultures per wound before the wound site was sealed with a transparent dressing (Tegaderm 3M, USA). At indicated time points, mice were euthanized and a 1 × 1 cm piece of skin encompassing the wound site was excised into 1 mL of 1X sterile PBS. Excised wounds were homogenized, and viable bacteria were enumerated by spotting onto respective selective agars. For OG1X and OG1RF selection, bacteria were spotted onto BHI solidified with 1.5% agar (Oxoid Technical No. 3) supplemented with 500 µg/mL streptomycin (MP Biomedicals, USA) or 25 µg/mL rifampicin (Sigma-Aldrich, USA), respectively. Animals that had lost the wound dressing at the time of sacrifice were excluded from data analysis. For competitive infection experiments, the competitive index (CI) was determined with the following formula:

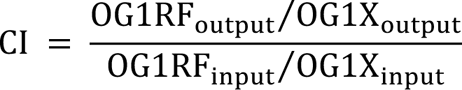

### Transposon sequencing

The *E. faecalis* transposon library containing ∼15,000 mutants was constructed and kindly provided to us by Gary M. Dunny (Kristich et al., 2008). Transposon mutants were arrayed into 96 wells with the transposon sequence of each mutant being known (Dale et al., 2018). Initial pools consisting of 100 transposon mutants per pool were made and glycerol stocked. Fifty transposon pools of 100 mutants were then combined to achieve a final pool size of 5,000. These pools were used for subsequent transposon sequencing, with a total of 3 distinct pools of 5,000 which covers the whole transposon library. Glycerol stocks of the 5,000 mutants were grown overnight in BHI medium (Neogen, Lansing, USA) for 15 - 18 h at 37 °C. Overnight cultures were washed twice with 1X sterile PBS after pelleting at 5,000 rpm for 4 min and normalized to OD_600_ of 0.35 corresponding to 2 × 10^8^ CFU/mL. Wounds were made on the dorsal back of the mice as described above and 10 μL of the normalized bacterial suspension was inoculated into wounds to achieve an infection CFU of 2 × 10^6^ CFU.

### Genomic DNA extraction for transposon sequencing

Mice were euthanized and wounds were excised at the indicated time points. Excised wounds were subsequently homogenized in 1 mL of 1X sterile PBS. To reduce biological variance, 500 μL from two wound homogenates containing the same transposon mutants were pooled together into 4 mL of BHI for a final volume of 5 mL at 37 °C for 3 h. The remaining wound homogenates were subjected to CFU enumeration on BHI agar (Acumedia, USA) supplemented with rifampicin or BHI agar without any antibiotics to check for the presence of contamination. Homogenates containing contaminants were excluded from subsequent library preparation. *In vitro* comparator pools were made by incubating 10 μL of normalized transposon overnight cultures into 4 mL of BHI broth at 37 °C for 3 h as well. To recover as many transposon mutants as possible, we included an enrichment step of transposon pools for all mice samples by incubating the mixture at 37 °C for 3 h, and DNA were extracted using the Qiagen DNeasy Blood and Tissue Kit (Qiagen, Germany). A total of 3 biological replicates of DNA samples were made from 6 wounds per transposon pool.

### Transposon library construction and sequencing

Extracted DNA was used for DNA library construction using NEBNext Ultra II DNA Library Prep Kit for Illumina (New England Biolabs, USA) according to the manufacturer’s instructions. DNA was subjected to acoustic shearing to obtain fragment sizes of approximately 300 bp using g-Tube (Covaris, USA). We adopted and modified the TraDIS protocol published by Barquist et al. (2016) by using the proposed splinkerette design for adapter ligation and subsequent enrichment of transposon pools using an amplicon-based sequencing approach. TraDIS adapters were used for adapter ligation and PCR amplified for final library construction using the Nextera XT DNA kit (Illumina, USA) as per the manufacturer’s instructions. Constructs were normalized and sequenced as 150 bp single read using the MiSeqV3. The sequencing was done by the sequencing facility at Singapore Centre for Environmental Life Sciences Engineering (SCELSE).

### Analysis of transposon sequencing results

Reads obtained from sequencing were checked using FastQC (version 0.11.5) and adapter trimmed using bbduk from the BBMap tools (version 34.49) (Bushnell, 2015). Trimmed reads containing the 15 bp transposon sequence at the 5’ region was obtained using a customized python script and subsequently trimmed to obtain sequences for mapping. Reads were mapped onto the *E. faecalis* OG1RF (NCBI accession: GCF_000172575.2) reference genome using BWA (version 0.7.15-r1140) (Li, H. & Durbin, 2009). Reads mapping to predicted open reading frames of each genome were quantified using HTSeq (Anders et al., 2015) and differential gene expression analysis was performed under the R environment (version 3.4.4) using Bioconductor package, edgeR (Robinson et al., 2010). Reads were normalized based on sequencing depth, scaled for the respective library sizes using trimmed mean of M-values (TMM), with common and tagwise dispersions being estimated for downstream analysis. Genes were considered differentially expressed with a false discovery rate (FDR) < 0.05 following correction by the Benjamin-Hochberg procedure. Gene annotation was performed using the database from the Kyoto Encyclopedia of Genes and Genomes (KEGG).

### RNA extraction from *in vivo* wound samples

A 1 × 1 cm of mouse skin encompassing the wound site was excised into 2 mL of RNA*later* stabilization solution (Invitrogen, USA) and incubated overnight at 4 °C. The mouse skin was then transferred into 1 mL of TRIzol (Life Technologies, USA) and cut into smaller pieces. The entire suspension was transferred into Lysing Matrix B 2 mL tubes (MP Biomedicals, USA) and homogenized using a FastPrep-24 tissue grinder (MP Biomedicals, USA) for 2 rounds of 40 s at 6.0 m/s with a 2 min rest on ice in between. To each sample, 200 µL of chloroform (Sigma-Aldrich, USA) was added, vortexed vigorously for 30 s and centrifuged at 12,000 × *g* for 10 min at 4 °C. The top layer (aqueous phase) containing RNA was transferred to 1.5 mL tubes containing 500 µL of ice-cold ethanol and shaken vigorously before loading into the RNeasy Mini spin columns (Qiagen, Germany). Subsequent RNA extraction steps were performed according to the manufacturer’s protocol of the RNeasy Mini Kit (Qiagen, Germany). Briefly, samples were washed once with Buffer RW1, followed by 2 washes with Buffer RPE and elution of RNA with RNase-free water. The RNA and potential DNA contamination concentrations were quantified using Qubit RNA BR and Qubit dsDNA HS assay kits, respectively. The extracted RNA was quality checked using the RNA ScreenTape on a TapeStation instrument (Agilent Technologies, USA). Every sample had a minimum RNA concentration of 100 ng/µL, a maximum of 10% DNA contamination and a RINe value ≥ 8.0 before it was used for library preparation. Library preparation was done using the Ribo-Zero Plus rRNA depletion kit (Illumina, USA) to remove mouse and bacterial rRNA from the extracted total RNA samples. The RNA samples were sequenced as 75 bp paired-end reads on an Illumina HiSeq2500. Library preparation and sequencing were carried out by the SCELSE sequencing facility.

### RNA extraction from *in vitro* bacterial cultures

Overnight cultures of *E. faecalis* wild-type OG1RF and OG1RF Δ*mptD* were sub-cultured to OD_600_ of 0.01 in a 24-well microtiter plate containing 1 mL of Tryptone Soya Broth without dextrose (TSBd, Sigma-Aldrich, USA) supplemented with or without mannose (Sigma-Aldrich, USA) in biological triplicates. Bacteria were harvested at late log/early stationary phase in RNAprotect Bacteria Reagent (Qiagen, Germany) and incubated at room temperature for 5 min before centrifugation at 10,000 *g* for 10 min. The supernatant was decanted, and bacteria pellets collected were subjected to total RNA extraction using the Qiagen RNeasy Mini Kit (Qiagen, Germany) with slight modifications. Briefly, cell pellets were resuspended in TE buffer containing 20 mg/mL lysozyme (Sigma-Aldrich, USA), further supplemented with 20 µL proteinase K (Qiagen, Germany), and incubated at 37 °C for 1 h. Subsequent RNA extraction steps were performed according to the manufacturer’s protocol. The extracted RNA samples were treated with DNase (TURBO DNA-*free* kit, Invitrogen, USA) for removal of genomic DNA before it was purified using the Monarch RNA cleanup kit (New England Biolabs, USA). The RNA and potential DNA contamination concentrations were quantified using Qubit RNA BR and Qubit dsDNA HS assay kits, respectively. The extracted RNA was quality checked using the RNA ScreenTape on a TapeStation instrument (Agilent Technologies, USA) before it was sent for sequencing. Every sample had a minimum RNA concentration of 40 ng/µL, a maximum of 10% DNA contamination and a RIN value ≥ 8.0 before being used for library preparation and subsequent sequencing as 75 bp paired-end reads on an Illumina HiSeq2500. Similarly, library preparation and sequencing were carried out by the SCELSE sequencing facility.

### *In vivo* and *in vitro* transcriptomic analysis

All raw reads obtained were checked using FastQC (Version 0.11.9) and adaptor trimmed using bbduk from BBMap tools (Version 39.79) (Bushnell, 2015). The trimmed reads were then mapped against *E. faecalis* OG1RF (NCBI accession: CP002621) reference genome using bwa-mem of BWA (Version 0.7.17-r1188) with default options. Reads mapped to open reading frames were quantified using htseq-count of HTSeq (Version 0.12.4) with option “-m intersection-strict” (Anders et al., 2015). All rRNA counts were manually removed from all data sets. Differential gene expression analysis was performed in R using edgeR (Version 3.28.1) (Robinson et al., 2010). The log2 fold change values extracted were based on the false discovery rate (FDR) ≤ 0.05. A gene set enrichment analysis (GSEA) was also done in R using the clusterProfiler package (Version 3.16.1) (Wu et al., 2021; Yu et al., 2012). For *in vitro* transcriptome analysis, common differentially expressed genes (DEGs) identified between without and with carbohydrate supplementation in TSBd media were removed, and only unique DEGs were used for downstream GSEA.

### Molecular cloning

The primers used in this study are listed in **Supplementary Table 2**. Transformants were screened using respective selective agar as follows: (1) *Escherichia coli* strains, LB with 500 µg/mL erythromycin (pGCP213 and pMSP3535) or 50 µg/mL kanamycin (pTCV); and (2) *E. faecalis* strains, BHI with 25 µg/mL erythromycin (pGCP213, pMSP3535 and pTCV). The generation of *E. faecalis* deletion mutants was done by allelic replacement using pGCP213 temperature-sensitive shuttle vector described previously (Nielsen et al., 2012). For the construction of OG1RF Δ*purEK* and OG1RF Δ*mptD*, vector pGCP213 was linearized using *Bam*HI and *Not*I (New England Biolabs, USA). Linearized pGCP213 and inserts were ligated using In-Fusion HD Cloning Kit (Clontech, Takara, Japan) and transformed into Stellar competent cells. Successful plasmid constructs were verified by Sanger sequencing and transformed into wild-type OG1RF. Transformants were selected with erythromycin at 30 °C, then passaged at non-permissive temperature at 42 °C with erythromycin to select for bacteria with successful plasmid integration into the chromosome. For plasmid excision, bacteria were serially passaged at 37 °C without erythromycin for erythromycin-sensitive colonies. These colonies were then subjected to PCR screening for detection of deletion mutants.

For the complementation of *purEK*, vector pTCV was linearized using *Bam*HI and *Sph*I (New England Biolabs, USA). Whereas, for the complementation of *mptD*, vector pMSP3535 was linearized using inverse PCR with iPCR_pMSP3535_F and iPCR_pMSP3535_R primers. Similarly, linearized pTCV and pMSP3535 as well as the respective inserts were ligated using In-Fusion HD Cloning Kit and transformed into Stellar competent cells. Successful plasmid constructs were verified by Sanger sequencing and transformed into OG1RF Δ*purEK* or OG1RF Δ*mptD*.

### Growth kinetic assay

Overnight cultures of the respective bacteria strains were diluted to OD_600_ of 0.01 in: (1) RPMI 1640 medium, no phenol red (Gibco, USA) supplemented with 1% (w/v) casamino acids (RPMI-CA; BD Biosciences, USA) with or without IMP; or (2) TSBd supplemented with 40 ng/mL nisin with or without 1% (w/v) galactose, mannose, or glucose (all purchased from Sigma-Aldrich, USA). A total of 200 μL of the diluted cell suspensions were inoculated per well in a 96-well polystyrene microtiter plate (Thermo Fisher Scientific, USA). Bacterial growth was measured at OD_600_ using a Tecan Infinite© M200 Pro spectrophotometer (Tecan Group Ltd, Switzerland) every 15 min for 16 h at 37 °C under static conditions.

### Purine metabolite quantification

Purine metabolites in mouse wounds were quantified using LC-MS performed by the Singapore Phenome Centre (SPC). A 1 × 1 cm of mouse skin encompassing the wound site was excised into 1.5 mL tubes, snap frozen in liquid nitrogen and submitted to the SPC for sample preparation, LC-MS profiling, and data processing. Briefly, tissue samples were weighed into tubes containing zirconium beads, 200 μL of 0.1M NaOH and 600 μL of methanol. The tubes were vortexed, homogenized and centrifuged at 10,000 rpm for 10 min at 4 °C. Two aliquots of the 300 μL supernatant were taken, dried down and reconstituted as follows: (1) for purine quantification, 100 μL of 80:20 acetonitrile:water, 15 mM of ammonium acetate, pH 9.2; and (2) for phosphate quantification, 100 μL of water, 20 mM of ammonium acetate, 0.1% formic acid.

Purine and phosphate quantification was performed on a Xevo TQ-S (Waters, UK). The source temperature was set at 150 °C with a cone gas flow of 150 L/h and a desolvation gas temperature of 450 °C with a desolvation gas flow of 900 L/h. The capillary voltage was set to 2.5 kV in the ESI positive or negative ionization mode for purine and phosphate quantification, respectively. For purine quantification, samples were injected into 2.1 mm × 100 mm, 1.7 μm UPLC BEH C18 column (Waters, UK) held at 45 °C. Mobile phase A is water with 15 mM of ammonium acetate (pH 9.2) and mobile phase B is 90:10 acetonitrile:water with 15 mM of ammonium acetate (pH 9.2). The column flow rate was 0.4 to 0.5 mL/min. For phosphate quantification, samples were injected into 2.1 mm × 150 mm, 1.7 μm UPLC HSS T3 column (Waters, UK) held at 45 °C. Mobile phase A is water with 20 mM of ammonium acetate and 0.1% formic acid. The column flow rate was 0.4 mL/min.

The weight of all tissue samples was measured prior to purine metabolite quantification, and the concentration of purine metabolites were normalized accordingly to the weight of the respective samples.

### Carbohydrate metabolism assay

Carbohydrate metabolism of *E. faecalis* strains were tested using API 50 CH (bioMérieux, France). Briefly, bacterial cultures were prepared to a turbidity equivalent of 2 McFarland standard and added to API 50 CHL Medium supplemented with 40 ng/mL nisin for induction of plasmid expression. The bacterial suspension was distributed into all 50 microtubes and sealed with mineral oil. The tray was then incubated at 37 °C under static condition for 48 h and 72 h with measurements taken at each time point to monitor for variability. The results for each microtube, positive (+), negative (-) and doubtful (?), were recorded.

### ELISA assay

A 1 × 1 cm of mouse skin encompassing the wound site was first rinsed in ice-cold 1X sterile PBS and homogenized in 1 mL of ice-cold 1X sterile PBS. The homogenates then undergo 2 freeze-thaw cycles to break the cell membranes and centrifuged at 5,000 × *g* for 5 min at 4 °C. The supernatants were collected and stored at −80 °C until assessment by Mouse Glucose ELISA, Mouse Galactose ELISA, and Mannose ELISA kits (MyBioSource, USA) as per the manufacturer’s protocol. Optical density of each well was determined at OD_450_ using a Tecan Infinite© M200 Pro spectrophotometer (Tecan Group Ltd, Switzerland).

### Urinary catheterization and bacterial infection

Bacterial cultures were normalized to 2 – 4 × 10^8^ CFU/mL in 1X sterile PBS. Implantation of catheters into mouse was performed as previously described (Guiton et al., 2010). Briefly, female C57BL/6 mice (8 – 9 weeks old, InVivos, Singapore) were anesthetized by inhalation of 3% isoflurane. Mice were inoculated with 50 μL of bacteria suspension (∼10^7^ CFU) into the urethra after catheterization. At 24 hpi, mice were euthanized. The bladders and kidneys were aseptically removed and homogenized in 1 mL and 800 μL of 1X sterile PBS, respectively. Catheters removed from the bladders were sonicated at 37 kHz and 30% power for 15 mins in 1 mL of 1X sterile PBS (Elma Ultrasonic, Germany), followed by vortexing at maximum speed for another 15 mins. All the samples were then serially diluted and viable bacteria were enumerated by spotting onto the respective selective agars. For OG1X and OG1RF selection, bacteria were spotted onto BHI agar supplemented with 500 µg/mL streptomycin (MP Biomedicals, USA) or 25 µg/mL rifampicin (Sigma-Aldrich, USA), respectively for competitive infection enumeration. Animals without catheters at the time of sacrifice were excluded from data analysis.

### Statistical analysis

Statistical analyses were performed with GraphPad Prism software (Version 9.0.0, California, USA) and are described in the respective figure legends.

### Ethics statement

All procedures were approved and performed in accordance with the Institutional Animal Care and Use Committee in Nanyang Technological University Singapore (ARF SBS/NIE-A19061, -A18063 and -A18059).

### Data availability

All the sequences have been deposited in the National Center for Biotechnology Information (NCBI) Gene Expression Omnibus (GEO) database under accession number GSE206751 (https://www.ncbi.nlm.nih.gov/geo/query/acc.cgi?acc=GSE206751).

## RESULTS

### *Eh. faecalis de novo* purine biosynthesis genes contribute to *E. faecalis* fitness during wound infection

Although *E. faecalis* is a common wound pathogen, little is known about the fitness determinants that contribute to wound infection. To identify genes contributing to fitness in wounds, using a mouse wound infection model that we have previously characterized for *E. faecalis* (Chong et al., 2017) and an *E. faecalis* OG1RF transposon mutant library consisting of ∼15,000 mutants (Kristich et al., 2008), we performed transposon sequencing (Tn-seq) at 8 h post-infection (hpi) and 3 days post-infection (dpi) following infection. Mutants disrupted in the *pur* operon (9 out of 11 genes) were significantly less abundant at 8 hpi compared to the pre-inoculation pool **(Table 1 and Supplementary Figure 1)**. The *E. faecalis pur* operon consists of 11 genes **(Supplementary Figure 2A)** and *de novo* purine biosynthesis undergoes 11 reactions from L-glutamine and 5-phosphoribosyl diphosphate (PRPP) to inosine monophosphate (IMP) before branching into specific pathways that produce guanosine and adenosine monophosphate (GMP and AMP) (Ramsey et al., 2014) **(Supplementary Figure 2B)**. At the same time point, we also performed *in vivo* RNA sequencing (RNA-seq) on *E. faecalis* wild-type OG1RF from infected wounds to provide a genome-wide analysis of differential gene expression during wound infection. We predicted that genes identified as essential for wound fitness using Tn-seq may display an increased gene expression, yielding a correlation coefficient between Tn-seq and RNA-seq close to -1. However, the calculated Spearman rank correlation coefficient between fold change of mutant abundance and fold change of differential expression among statistically significant genes was −0.0311 **(Supplementary Figure 3A)**, which was similar to a previous study looking at fitness determinants and gene expression during *P. aeruginosa* wound infection (Turner et al., 2014). Nevertheless, our observation that all 11 genes in the *pur* operon were significantly upregulated at 8 hpi compared to inoculum **(Table 2 and Supplementary Figure 3B)**, suggested a role for purine biosynthesis during acute *E. faecalis* wound infection.

**Table 1.** E. faecalis transposon mutant abundance profiled by Tn-seq from 8 hpi wounds.

| Locus tag | Name | Description | log<sub>2</sub>FC | p-value | FDR |
| --- | --- | --- | --- | --- | --- |
| OG1RF_10019 | *mptAB* | PTS mannose transporter subunit EIIAB | -1.15 | 5.75E-06 | 2.11E-05 |
| OG1RF_10020 | *mptC* | PTS mannose/fructose/sorbose transporter subunit IIC | -0.92 | 3.54E-10 | 1.58E-09 |
| OG1RF_10021 | *mptD* | PTS mannose transporter subunit IID | -1.11 | 8.90E-18 | 7.46E-17 |
| OG1RF_11489 | *purD* | phosphoribosylamine--glycine ligase | -1.70 | 1.12E-06 | 3.48E-06 |
| OG1RF_11490 | *purH* | inosine monophosphate cyclohydrolase | -2.29 | 1.61E-37 | 5.62E-36 |
| OG1RF_11491 | *purN* | phosphoribosylglycinamide formyltransferase | -1.16 | 1.63E-02 | 3.03E-02 |
| OG1RF_11492 | *purM* | phosphoribosylformylglycinamidine cyclo-ligase | -2.06 | 1.87E-15 | 1.30E-14 |
| OG1RF_11493 | *purF* | amidophosphoribosyltransferase | -3.19 | 3.23E-27 | 1.82E-26 |
| OG1RF_11494 | *purL* | phosphoribosylformylglycinamidine synthase II | -1.79 | 1.38E-11 | 3.68E-11 |
| OG1RF_11495 | *purL2* | phosphoribosylformylglycinamidine synthase subunit PurQ | -3.03 | 2.91E-37 | 2.59E-36 |
| OG1RF_11496 | *purS* | phosphoribosylformylglycinamidine synthase | -2.68 | 7.75E-15 | 5.12E-14 |
| OG1RF_11498 | *purK* | 5-(carboxyamino)imidazole ribonucleotide synthase | -2.15 | 9.76E-13 | 5.37E-12 |

Complete table can be found in Supplementary file 1 (sheet 1).

**Table 2.** E. faecalis purine biosynthesis genes differentially regulated from 8 hpi wounds.

| Locus tag | Name | Description | log<sub>2</sub>FC | p-value | FDR |
| --- | --- | --- | --- | --- | --- |
| OG1RF_11489 | *purD* | phosphoribosylamine--glycine ligase | 6.47 | 5.40E-199 | 4.36E-197 |
| OG1RF_11490 | *purH* | inosine monophosphate cyclohydrolase | 6.26 | 2.78E-182 | 1.50E-180 |
| OG1RF_11491 | *purN* | phosphoribosylglycinamide formyltransferase | 6.27 | 1.91E-115 | 3.48E-114 |
| OG1RF_11492 | *purM* | phosphoribosylformylglycinamidine cyclo-ligase | 7.02 | 7.40E-132 | 1.73E-130 |
| OG1RF_11493 | *purF* | amidophosphoribosyltransferase | 7.40 | 1.57E-141 | 4.15E-140 |
| OG1RF_11494 | *purL* | phosphoribosylformylglycinamidine synthase II | 6.58 | 9.95E-152 | 3.10E-150 |
| OG1RF_11495 | *purL2* | phosphoribosylformylglycinamidine synthase subunit PurQ | 6.12 | 1.16E-82 | 1.35E-81 |
| OG1RF_11496 | *purS* | phosphoribosylformylglycinamidine synthase | 6.35 | 7.15E-22 | 2.25E-21 |
| OG1RF_11497 | *purC* | phosphoribosylaminoimidazolesuccinocarboxamide synthase | 7.29 | 6.75E-118 | 1.28E-116 |
| OG1RF_11498 | *purK* | 5-(carboxyamino)imidazole ribonucleotide synthase | 5.37 | 1.57E-62 | 1.20E-61 |
| OG1RF_11499 | *purE* | 5-(carboxyamino)imidazole ribonucleotide mutase | 5.24 | 1.24E-75 | 1.29E-74 |

*Complete table can be found in Supplementary file 1 (sheet 2).*

To confirm the role of *de novo* purine biosynthesis for *E. faecalis* replication during wound infection, we created an in-frame deletion mutant of the first two genes in the *pur* operon, *purEK* (OG1RF Δ*purEK*) (Kilstrup et al., 2005) and performed an *in vivo* competitive infection with *E. faecalis* OG1X at 8 hpi, a closely related strain expressing different antibiotic resistance genes enabling differential selection (Ike et al., 1983). In accordance with our Tn-seq and RNA-seq results, OG1RF Δ*purEK* (CI = 0.93) displayed significantly reduced fitness compared to wild-type OG1RF (CI = 1.69) **(Figure 1A)**. We subsequently performed single-strain infection and observed that OG1RF Δ*purEK* CFU (8.33 × 10^6^ CFU/wound) were significantly lower than wild-type OG1RF (2.77 × 10^7^ CFU/wound) at 8 hpi **(Figure 1B)**.

**Figure 1.**
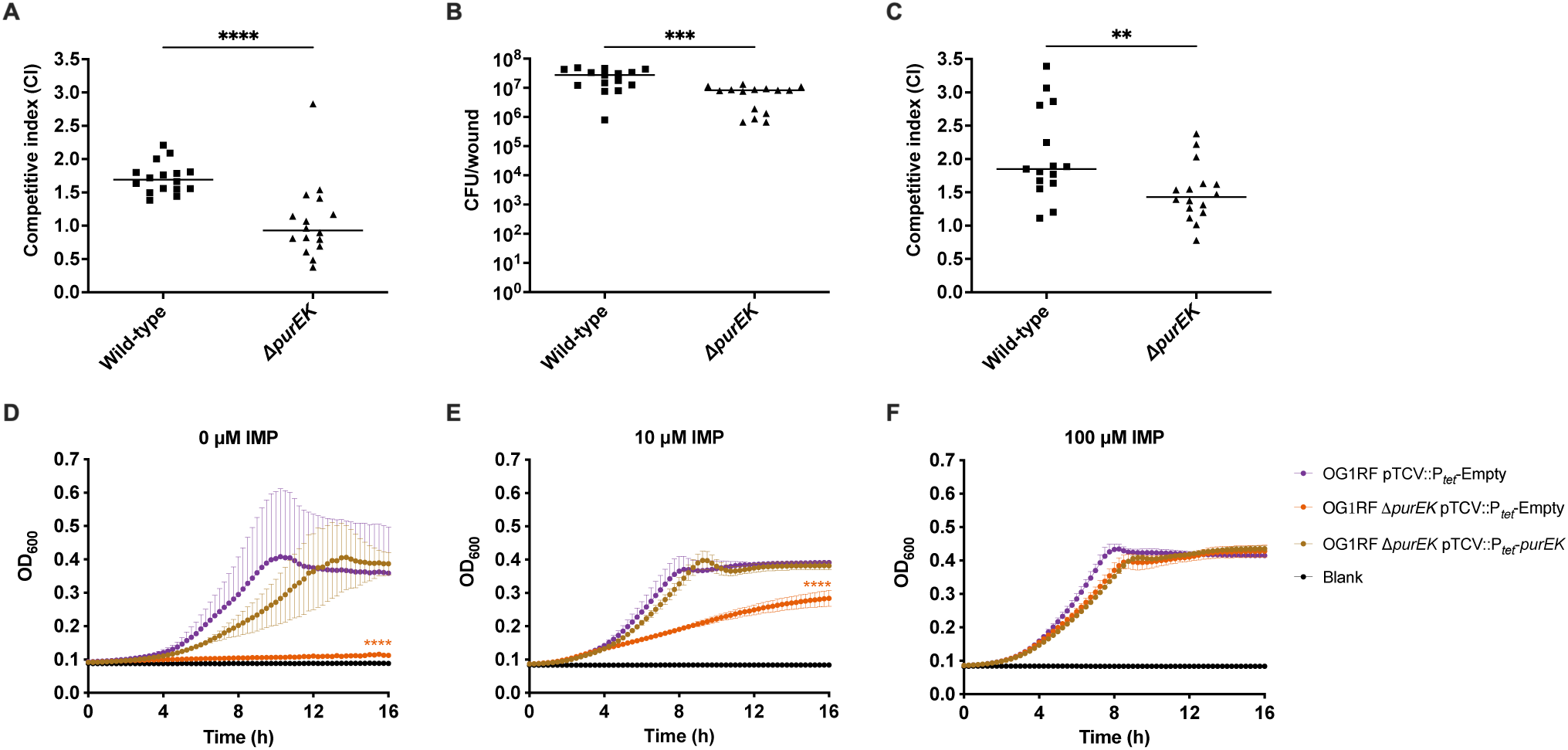
*De novo* purine biosynthesis contributes to *E. faecalis* fitness during early stages of wound infection. Male C57BL/6 mice were wounded and infected with **(A)** a 1:1 ratio of *E. faecalis* OG1X:wild-type OG1RF or OG1X:OG1RF Δ*purEK* at 2 – 4 × 10^6^ CFU/wound, **(B)** 2 – 4 × 10^6^ CFU of wild-type OG1RF or OG1RF Δ*purEK* in single-strain infection and CFU determined at 8 hpi, or **(C)** a 1:1 ratio of *E. faecalis* OG1X:wild-type OG1RF or OG1X:OG1RF Δ*purEK* at 2 – 4 x 10^6^ CFU/wound and CFU determined at 3 dpi. The recovered bacteria were enumerated on selective agar plates for each strain. Each data point represents one mouse and horizontal lines indicate the median, N = 3, n = 5 - 6 mice per group per experiment. Statistical analysis was performed using the Mann-Whitney U test, **p < 0.01, ***p < 0.001, ****p < 0.0001. Growth kinetics of wild-type OG1RF pTCV::P*_tet_*-Empty, OG1RF Δ*purEK* pTCV::P*_tet_*-Empty, and OG1RF Δ*purEK* pTCV::P*_tet_*-*purEK* in RPMI-CA media supplemented with **(D)** 0, **(E)** 1 or **(F)** 100 μM IMP over 16 h. Baseline readings are indicated by Blank, containing only the growth media. Data are mean values of three independent biological replicates and vertical lines represent SD from the mean. Statistical analysis was performed at 16 h OD_600_ measurement with wild-type OG1RF pTCV::P*_tet_*-Empty as the comparator using the Mann-Whitney U test, ****p < 0.0001.

Tn-seq analysis at 3 dpi also suggested the importance of *de novo* purine biosynthesis for persistence in wounds, as two genes in the *pur* operon (*purH* and *purM*) were significantly less abundant in the post-infection transposon pools **(Table 3)**. To validate the contribution of purine biosynthesis at 3 dpi, we similarly performed the *in vivo* competitive infection with OG1X. Although OG1RF Δ*purEK* (CI = 1.43) displayed significantly reduced fitness compared to wild-type OG1RF (CI = 1.85) at 3 dpi **(Figure 1C)**, the difference in CI was not as large as compared to 8 hpi **(Figure 1A)**. Overall, these results demonstrate that *E. faecalis de novo* purine biosynthesis contributes significantly to *E. faecalis* replication during acute infection as well as to persistence during wound infection.

**Table 3.**
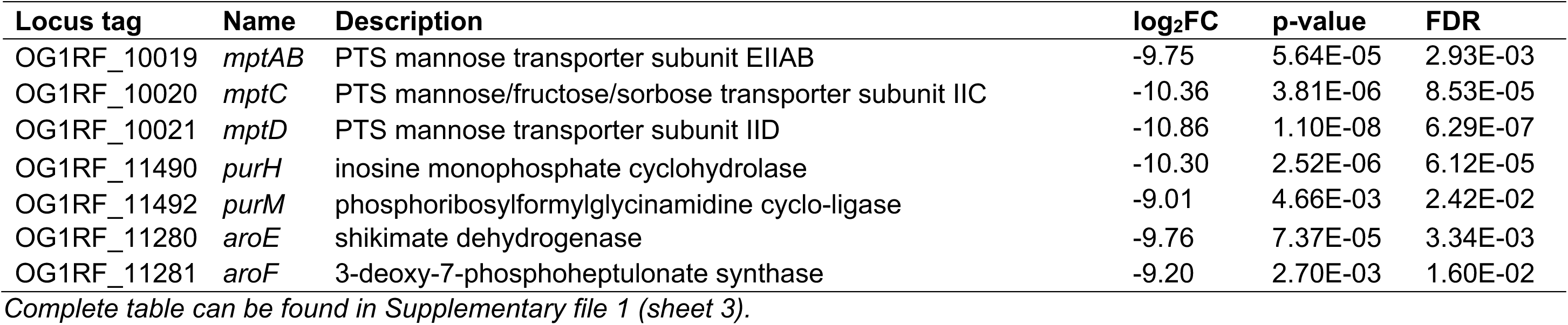
*E. faecalis* transposon mutant abundance profiled by Tn-seq from 3 dpi wounds.

We next validated the predicted requirement of *de novo* purine biosynthesis for *E. faecalis* growth in RPMI-CA medium lacking purines. As expected, the deletion of *purEK* (OG1RF Δ*purEK* pTCV::P*_tet_*-Empty) resulted in severe growth attenuation compared to wild-type (OG1RF pTCV::P*_tet_*-Empty), and complementation of *purEK* on a plasmid (OG1RF Δ*purEK* pTCV::P*_tet_*-*purEK*) restored growth to near wild-type levels **(Figure 1D)**. To confirm that the disruption of the purine biosynthesis pathway was the sole reason for the growth attenuation observed, we supplemented the RPMI-CA medium with 10 and 100 μM IMP (end-product of purine biosynthesis; **Supplementary Figure 2B**). With supplementation of IMP, the growth of OG1RF Δ*purEK* pTCV::P*_tet_*-Empty was restored to wild-type levels in a dose-dependent manner **(Figure 1E and 1F)**. These findings demonstrate the importance of the *pur* operon for *E. faecalis* growth in a purine-deficient environment.

### Purine metabolites in wounds are low during the acute replication phase of *E. faecalis* infection

*De novo* purine biosynthesis is required for *Staphylococcus aureus* pathogenesis during bacteremia because the disruption of purine biosynthesis leads to reduced bacterial virulence as measured by animal weight loss and bacterial burden (Goncheva et al., 2020). Moreover, previous Tn-seq studies revealed that purines were among the metabolites deemed “not available” to *P. aeruginosa* during both burn and chronic wound infections (Turner et al., 2014). Collectively, these studies demonstrate the significance of purine biosynthesis for virulence across various bacteria and infection sites. Hence, we predicted that purine availability in the wound microenvironment is low, which would explain the importance of *de novo* purine biosynthesis for successful *E. faecalis* wound infection. To assess whether purines were indeed limited during wound infection, we quantified purine metabolites from *E. faecalis* wild-type OG1RF-infected wounds compared to PBS-treated control wounds at 8 hpi and 3 dpi using LC-MS. Following infection with *E. faecalis* wild-type, we observed a trend towards decreased purine metabolites compared to PBS-treated wounds at 8 hpi which was significant for adenine, guanine, xanthine, and AMP **(Figure 2A – 2D**, compare blue bars), suggesting that there may be consumption of purine metabolites during acute wound infection which could explain the high demand for purine metabolites when *E. faecalis* is rapidly replicating in the wounds. By contrast, the levels of purine metabolites remain similar between PBS-treated and wild-type OG1RF-infected wounds at 3 dpi, except for xanthine and AMP **(Figure 2C and 2D**, compare green bars). Differences following infection were not observed for adenosine, guanosine, and inosine **(Supplementary Figure 4)**. Taken together, our results suggest that the importance of *de novo* purine biosynthesis for *E. faecalis* replication during acute wound infection is driven by its rapid consumption of purine metabolites in the wound microenvironment.

**Figure 2.**
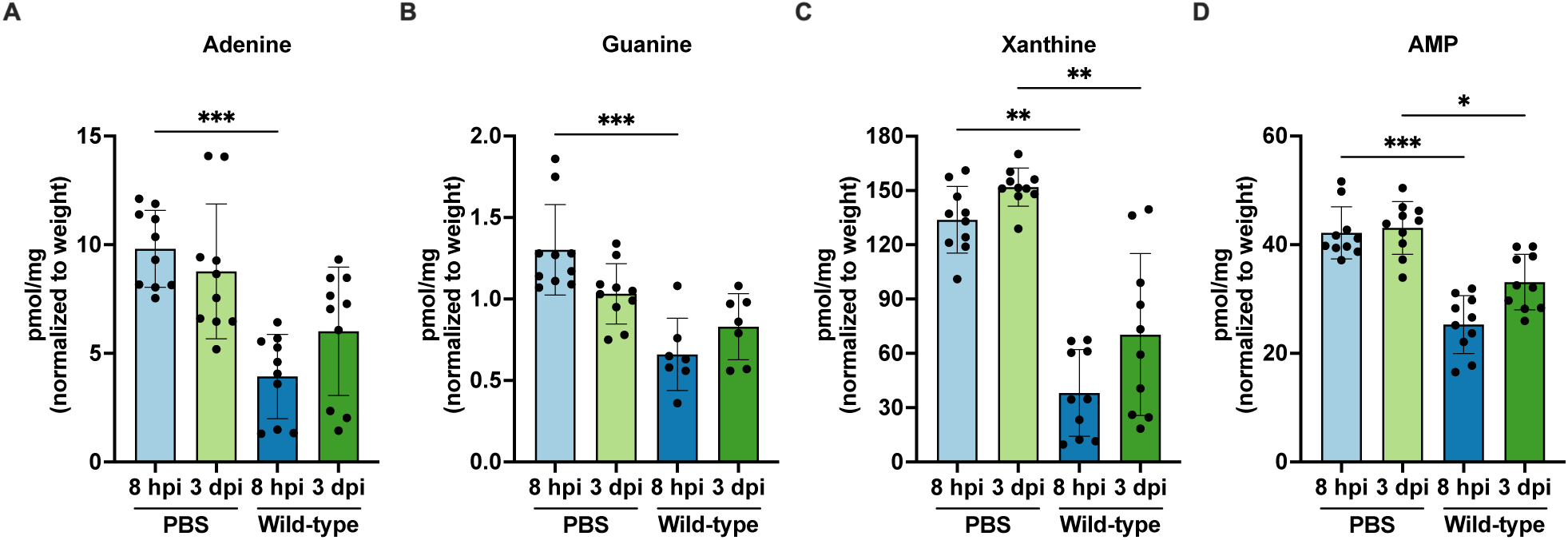
Purine metabolites are low during early *E. faecalis* wound infection. Male C57BL/6 mice were wounded and inoculated with PBS or 2 – 4 × 10^6^ CFU of wild-type OG1RF. Wounds were harvested at 8 hpi and 3 dpi for quantification of **(A)** adenine, **(B)** guanine, **(C)** xanthine, and **(D)** AMP using LC-MS. Each data point represents one mouse and error bars represent SD from the mean; N = 2, n = 5 mice per group per experiment. Statistical analysis was performed using the Mann-Whitney U test, *p < 0.05, **p < 0.01, ***p < 0.001.

### *E. faecalis* MptABCD phosphotransferase system is important for *E. faecalis* persistence during wound infection

Tn-seq analysis at 8 hpi and 3 dpi also identified *mptABCD* as significantly contributing to *E. faecalis* fitness at both time points **(Tables 1 and 3)**. We therefore investigated the contribution of *mptABCD* to *E. faecalis* virulence during wound infection, especially at 3 dpi when transposon insertions in all genes of the *mpt* operon were among the most significantly underrepresented following Tn-seq **(Supplementary Figure 5)**. *E. faecalis mptABCD* encodes a carbohydrate-specific phosphotransferase system used for the import of carbohydrates (Deutscher et al., 2006) to facilitate wound persistence.

To validate the role of the *E. faecalis* MptABCD PTS during wound infection, we created an in-frame deletion mutant of *mptD* (OG1RF Δ*mptD*) and performed an *in vivo* competitive infection with OG1X at 8 hpi and 3 dpi. In agreement with our Tn-seq results, OG1RF Δ*mptD* had reduced fitness compared to wild-type OG1RF at both 8 hpi and 3 dpi **(Figure 3A and 3B)**. However, we observed a bigger difference in CI at 3 dpi (CI_OG1RF_ = 1.89, CI_Δ*mptD*_ = 0.23) compared to 8 hpi (CI_OG1RF_ = 1.53, CI_Δ*mptD*_ = 1.08), which may explain the more pronounced decrease in log_2_FC that we observed in the post-infection transposon pools at 3 dpi **(Tables 1 and 3)**. Similarly, we performed single-strain infection and observed that OG1RF Δ*mptD* (3.50 × 10^4^ CFU/wound) colonized poorer than wild-type OG1RF (2.47 × 10^5^ CFU/wound) at 3 dpi **(Figure 3C)**. Together, these results indicate that MptABCD PTS plays a significant role during *E. faecalis* persistence in wounds.

**Figure 3.**
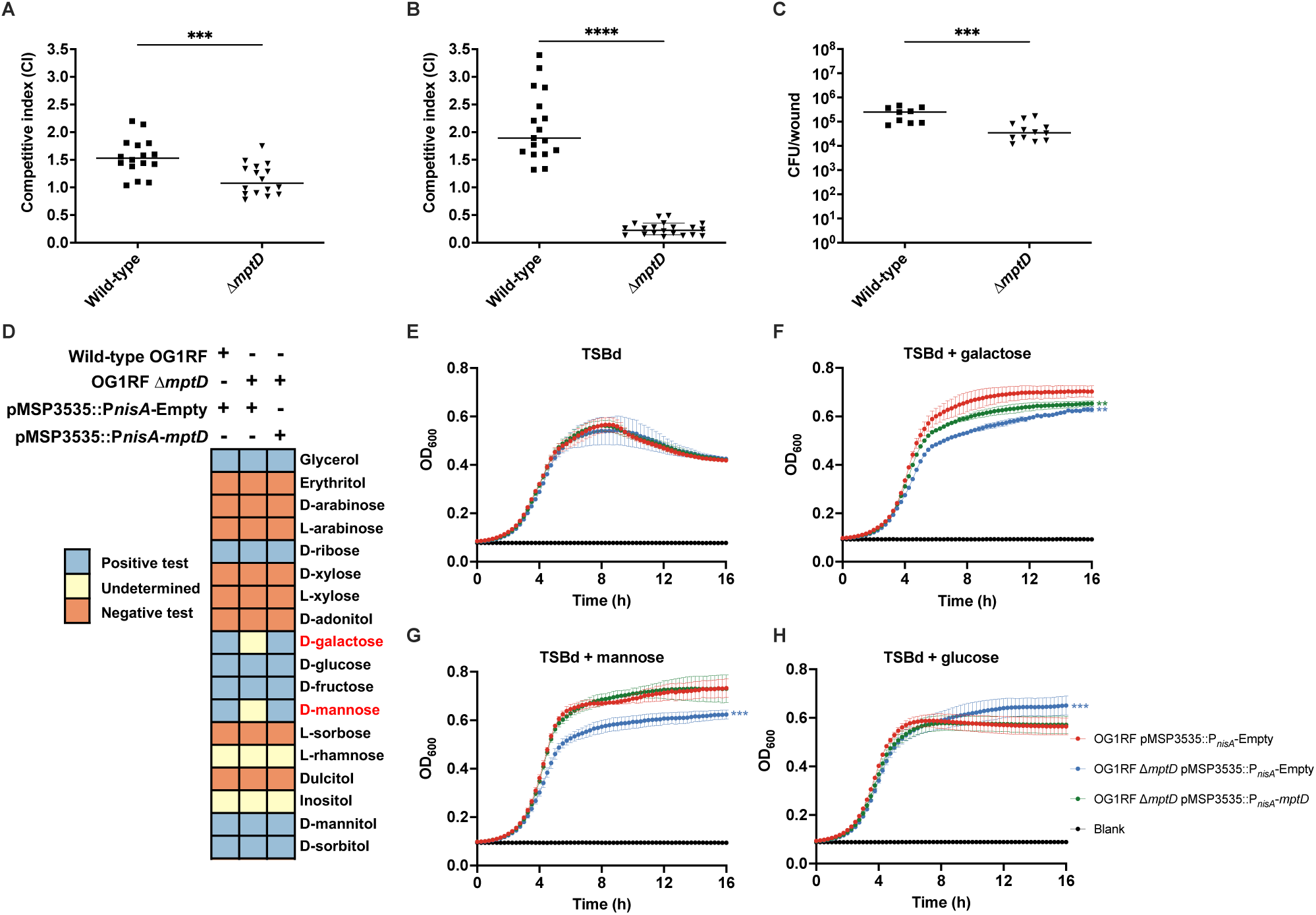
MptABCD phosphotransferase system contributes to *E. faecalis* wound fitness during persistence. Male C57BL/6 mice were wounded and infected with **(A)** a 1:1 ratio of *E. faecalis* OG1X:wild-type OG1RF or OG1X:OG1RF Δ*mptD* at 2 – 4 × 10^6^ CFU/wound (N = 3, n = 5 - 6 mice) and CFU determined at 8 hpi or **(B)** a 1:1 ratio of *E. faecalis* OG1X:wild-type OG1RF or OG1X:OG1RF Δ*mptD* at 2 – 4 × 10^6^ CFU/wound (N = 4, n = 5 - 6 mice) and CFU determined at 3 dpi or **(C)** 2 – 4 × 10^6^ CFU of wild-type OG1RF or OG1RF Δ*mptD* (N = 2, n = 5 - 6 mice) and CFU determined at 3 dpi. The recovered bacteria were enumerated on selective agar plates for each strain. Each data point represents one mouse and horizontal lines indicate the median. Statistical analysis was performed using the Mann-Whitney U test, ***p < 0.001, ****p < 0.0001. **(D)** Carbohydrate fermentation test (API 50 CH) of wild-type OG1RF pMPSP3535::P*_nisA_*-Empty, OG1RF Δ*mptD* pMSP3535::P*_nisA_*-Empty, and OG1RF Δ*mptD* pMSP3535::*P_nisA_-mptD*. Plasmid-based *mptD* expression was induced with 40 ng/mL nisin. Results shown are a subset of the 50 carbohydrates; differences were only detected for D-galactose and D-mannose (see **Supplementary Table 3** for complete table). Positive tests were determined by a change of the bromcresol purple indicator in the medium to yellow. For undetermined test results, the bromcresol purple indicator did not change to yellow nor did remain purple. Negative tests occurred when the bromcresol purple indicator remained purple. Growth kinetics of wild-type OG1RF pMPSP3535::P*_nisA_*-Empty, OG1RF Δ*mptD* pMSP3535::P*_nisA_*-Empty, and OG1RF Δ*mptD* pMSP3535::*P_nisA_-mptD* in TSBd media supplemented **(E)** without additional carbohydrates and with 1% (w/v) **(F)** galactose, **(G)** mannose, and **(H)** glucose over 16 h. Plasmid-based *mptD* expression was induced with 40 ng/mL nisin. Baseline readings are indicated by Blank, containing only the growth media. Data are mean values of three independent biological replicates and vertical lines represent SD from the mean. Statistical analysis was performed at 16 h OD_600_ measurement with wild-type OG1RF pMPSP3535::P*_nisA_*-Empty as the comparator using the Mann-Whitney U test, **p < 0.01, ***p < 0.001.

### *E. faecalis* MptABCD phosphotransferase system is responsible for the import of galactose and mannose

Based on KEGG genome, *E. faecalis mptABCD* is predicted to encode PTS mannose/fructose/sorbose transporter subunits. Therefore, to determine the carbohydrate(s) transported by the MptABCD PTS, we tested the ability of *E. faecalis* to metabolize 50 different carbohydrates. Across all 50 carbohydrates, the deletion of *mptD* (OG1RF Δ*mptD* pMSP3535::P*_nisA_*-Empty) only affected the metabolism of galactose and mannose when compared to wild-type (OG1RF pMSP3535::P*_nisA_*-Empty), and complementation of *mptD* on an inducible plasmid (OG1RF Δ*mptD* pMSP3535::P*_nisA_*-*mptD*) restored the mannose and galactose metabolism **(Figure 3D)**. To validate that the *mptD* deletion mutant was indeed unable to metabolize galactose and mannose, we performed growth kinetic assays of wild-type OG1RF, *mptD* deletion, and complement strains in TSBd (TSB broth lacking dextrose) growth media supplemented with different carbohydrates. We did not observe any growth differences between all three strains in TSBd in the absence of carbohydrate supplementation **(Figure 3E)**. When TSBd was supplemented with either galactose or mannose, the growth of OG1RF pMSP3535::P*_nisA_*-Empty was augmented, but OG1RF Δ*mptD* pMSP3535::P*_nisA_*-Empty growth was not **(Figure 3F and 3G)**. Complementation of *mptD* in the deletion mutant resulted in significantly improved growth compared to wild-type OG1RF pMSP3535::P*_nisA_*-Empty levels when supplemented with galactose **(Figure 3F)** and was almost identical to wild-type OG1RF pMSP3535::P*_nisA_*-Empty levels with mannose supplementation **(Figure 3G)**. As a control, we also supplemented TSBd with glucose, for which we did not expect to observe any growth differences between the three strains as metabolism of glucose was unaffected when *mptD* was deleted **(Figure 3D)**. As expected, growth differences were similar between all strains, at least through log phase, when supplemented with glucose **(Figure 3H)**. Collectively, these results indicate that MptABCD PTS is responsible for the import for galactose and mannose into *E. faecalis*, and that the import of these carbohydrates may contribute to *E. faecalis* persistence in wounds.

### Carbohydrate availability changes as the wound infection progresses

Galactose and mannose import appear to be more crucial at 3 dpi than at 8 hpi (**Figure 3A and 3B)**. We reasoned that carbohydrate availability in wounds may change as wound infection progresses, where successful *E. faecalis* persistence is dependent on its promiscuous ability to source from a wide array of nutrients in the wound microenvironment. We hypothesized that these changing carbohydrate availability influences the natural course of wound pathogenesis, as well as *E. faecalis* depletion of preferred carbohydrate sources such as glucose during acute infection, would necessitate a switch to other carbohydrates such as mannose and galactose at later time points. To test this hypothesis, we harvested PBS-treated, *E. faecalis* wild-type OG1RF- and OG1RF Δ*mptD*-infected wounds at 8 hpi and 3 dpi, and quantified the concentrations of glucose, galactose, and mannose. Consistent with our prediction, we detected lower glucose concentrations at 3 dpi compared to 8 hpi in PBS-treated wounds **(Figure 4A)**, and concordant higher galactose and mannose concentrations at 3 dpi compared to 8 hpi **(Figure 4B and 4C)**. Additionally, *E. faecalis* wild-type OG1RF infection further decreased glucose, galactose and mannose concentrations compared to PBS-treated wounds at 8 hpi **(Figure 4A – 4C)**, suggesting that *E. faecalis* can use all three carbohydrates during early phases of wound infection, which may support its rapid growth. At 3 dpi however, we detected no significant differences in glucose concentrations in any infected wounds **(Figure 4A)** and observed that *E. faecalis* wild-type OG1RF- and OG1RF Δ*mptD*-infected wounds had lesser galactose than PBS-treated wounds **(Figure 4B)**. Likewise, lesser mannose was detected compared to PBS-treated wounds when infected with *E. faecalis* wild-type OG1RF at 3 dpi **(Figure 4C)**. These results indicate that *E. faecalis* can import and deplete galactose and mannose availability during wound infection. Consistent with this conclusion, mannose concentrations were similar between PBS-treated and OG1RF Δ*mptD*-infected wounds at 8 hpi and higher in OG1RF Δ*mptD*-infected wounds compared to OG1RF-infected wounds at 3 dpi (albeit not significant) **(Figure 4C)**, suggesting that disruption of MptABCD PTS indeed leads to decreased mannose import (i.e. mannose accumulation) *in vivo*. Unexpectedly, we did not detect significant differences in galactose concentration between wild-type OG1RF- and OG1RF Δ*mptD*-infected wounds at any time point **(Figure 4B)**, despite a role for MptABCD in galactose metabolism *in vitro* **(Figure 3D)**. Since we observed minimal glucose depletion by *E. faecalis* at 3 dpi, we wondered whether a high glucose microenvironment (such as that found during hyperglycemia in diabetic mice) would prompt *E. faecalis* to import glucose instead of galactose and mannose, and consequently render the MptABCD PTS dispensable during persistence in diabetic animals. As such, we performed an *in vivo* competitive infection of OG1RF Δ*mptD* with OG1X in diabetic (*db/db*) mice at 3 dpi. However, like non-diabetic mice, OG1RF Δ*mptD* (CI = 0.34) had reduced fitness compared to wild-type OG1RF (CI = 1.22) at 3 dpi **(Supplementary Figure 6)**. Taken together, these results suggest that as the wound infection progresses, glucose becomes depleted and other carbohydrates, such as galactose and mannose, become more available in the wounds. As such, *E. faecalis* undergo a metabolic switch towards galactose and mannose metabolism as the wound infection progresses.

**Figure 4.**
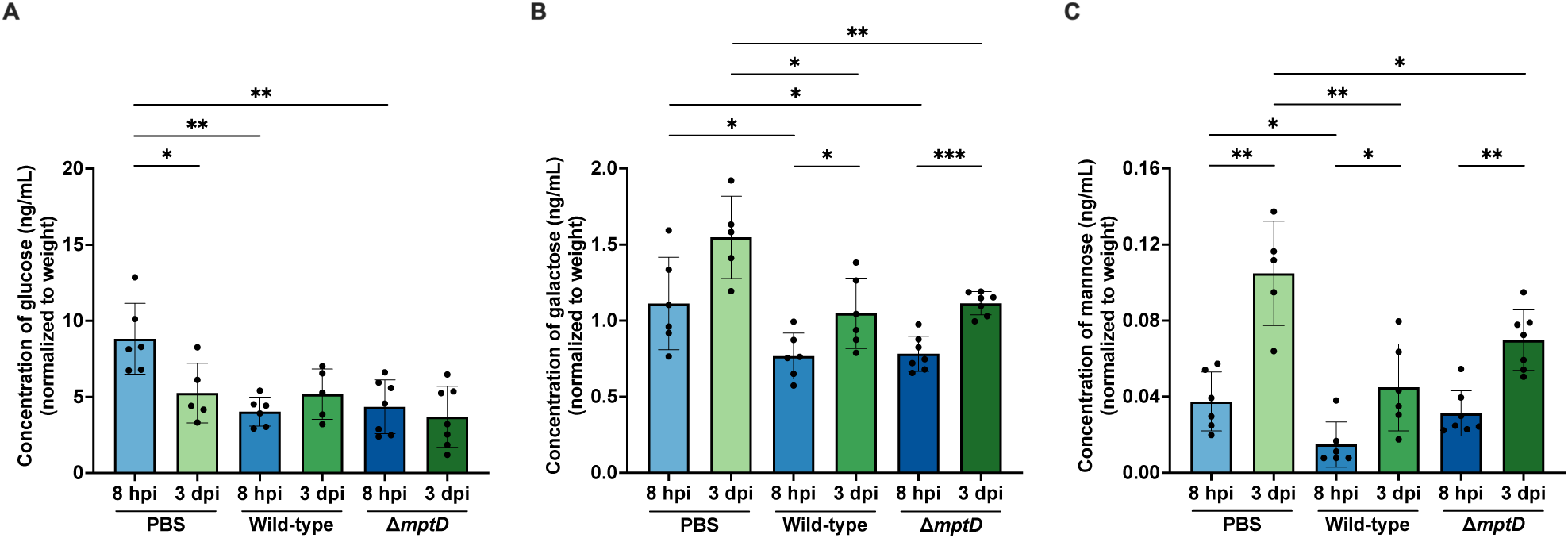
Carbohydrate availability changes as *E. faecalis* wound infection progresses. Male C57BL/6 mice were wounded and inoculated with sterile PBS, wild-type OG1RF or OG1RF Δ*mptD* at 2 – 4 × 10^6^ CFU/wound. Wounds were harvested at 8 hpi and 3 dpi, and subjected to **(A)** glucose, **(B)** galactose and **(C)** mannose quantification by ELISA. Each data point represents measurement from one mouse; error bars represent SD from the mean; N = 1, n = 6 - 7 mice. Statistical analysis was performed using the Mann-Whitney U test, *p < 0.05, **p < 0.01, ***p < 0.001.

### *E. faecalis de novo* purine and shikimate biosynthesis are regulated by MptABCD phosphotransferase system-mediated mannose import

Since MptABCD PTS imports galactose and mannose and this contributes to *E. faecalis* virulence *in vivo*, we asked whether there were any galactose and/or mannose-dependent changes in gene expression that might further explain why in particular these carbohydrates are important during wound infection. We therefore performed *in vitro* RNA-seq with wild-type OG1RF and OG1RF Δ*mptD* grown in TSBd without and with supplementation of galactose or mannose. Based on gene set enrichment analysis (GSEA), we did not observe any gene sets enriched with the supplementation of galactose. By contrast, when wild-type OG1RF and OG1RF Δ*mptD* were grown in TSBd supplemented with mannose, we observed several processes/pathways such as purine metabolism, PTS, fructose and mannose metabolism, biosynthesis of secondary metabolites and amino acids that were enriched **(Figure 5A)**. Among the enriched processes/pathways, 7 out of 8 genes in the shikimate biosynthesis operon (*aroF*, *aroE*, *aroC*, *tyrA*, *aroA*, *aroK*, and *pheA*) and 6 out of 11 genes in the *pur* operon (*purH*, *purN*, *purM*, *purF*, *purL* and *purL2*) were significantly downregulated in OG1RF Δ*mptD* when grown in TSBd supplemented with mannose **(Figure 5A)**, suggesting that shikimate and purine biosynthesis are attenuated when import of mannose is hindered. KEGG pathway analysis showed that mannose imported by MptABCD PTS was functionally linked to shikimate and purine biosynthesis **(Figure 5B)**. Following mannose import by MptABCD PTS, it can undergo a series of reactions that leads to the production of PEP (a substrate of shikimate pathway) or production of PRPP (a substrate for purine biosynthesis) **(Figure 5B)**. These results were also supported by Tn-seq analysis whereby transposon mutants for some genes of the shikimate biosynthesis and *pur* operon were significantly underrepresented in the post-infection transposon pools at 3 dpi **(Table 3)**.

**Figure 5.**
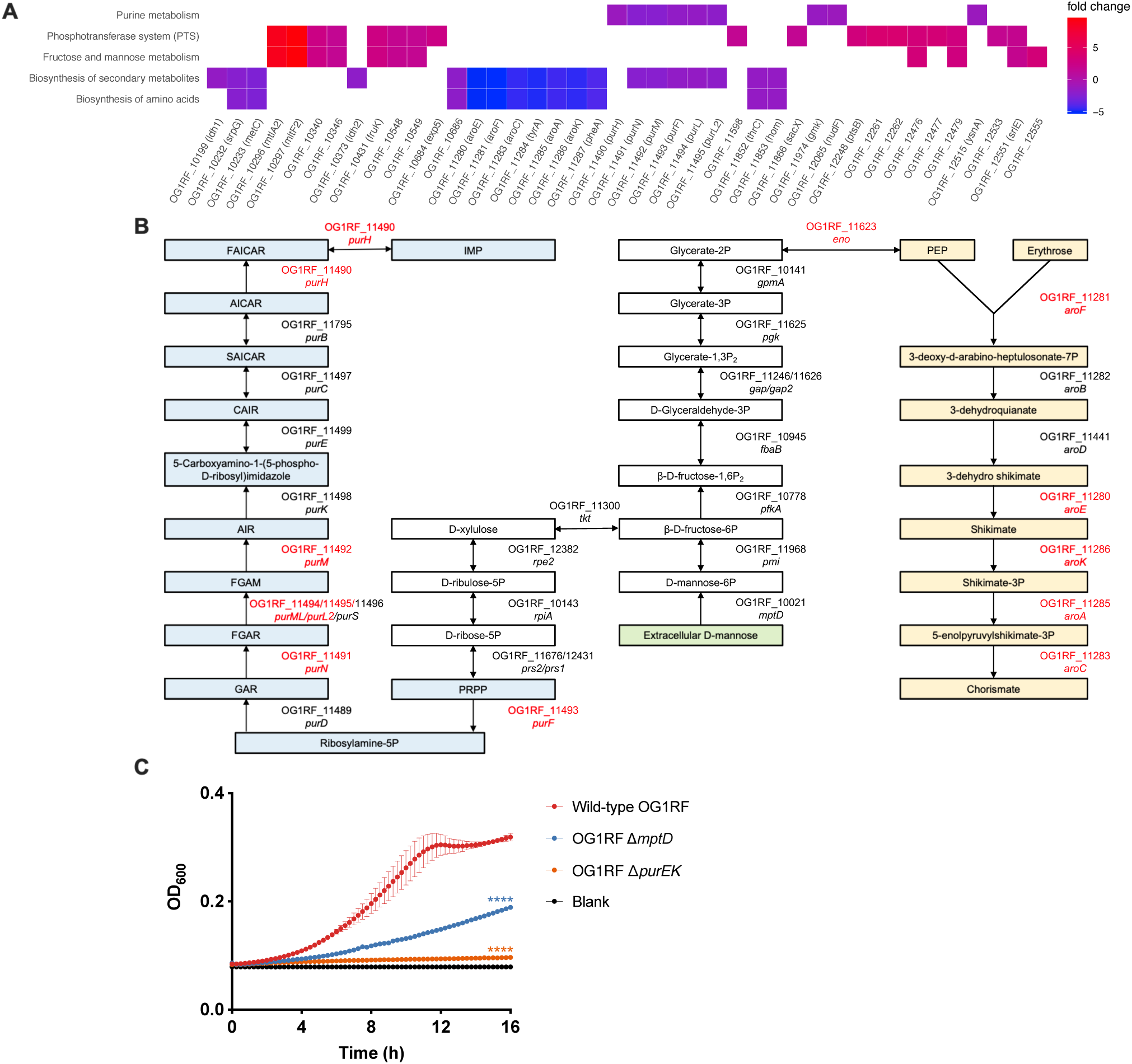
Mannose imported by *E. faecalis mptABCD* is functionally linked to *de novo* purine and shikimate biosynthesis. **(A)** Gene set enrichment pathways identified based on differentially expressed genes between wild-type OG1RF and OG1RF Δ*mptD* grown in TSBd supplemented with 1% (w/v) mannose. Complete table of differentially expressed genes can be found in Supplementary file 1 (sheet 4 – 8). **(B)** KEGG pathways depicting how import of extracellular D-mannose by MptD (green) acts as a substrate for *de novo* purine (blue) and shikimate (yellow) biosynthesis. **(C)** Growth kinetics of wild-type OG1RF, OG1RF Δ*mptD*, and OG1RF Δ*purEK* in RPMI-CA over 16 h. Baseline readings are indicated by Blank, containing only the growth media. Data are mean values of three independent biological replicates and vertical lines represent SD from the mean. Statistical analysis was performed at 16 h OD_600_ measurement with wild-type OG1RF as the comparator using the Mann-Whitney U test, ****p < 0.0001.

To subsequently validate that purine biosynthesis was impeded when mannose import was hindered in OG1RF Δ*mptD*, we performed a growth kinetic assay in RPMI-CA medium lacking purines. In agreement with our *in vitro* RNA-seq analysis, the growth of OG1RF Δ*mptD* was attenuated compared to wild-type OG1RF in RPMI-CA medium **(Figure 5C)**. These results confirm that hindered mannose import by MptABCD PTS also reduces *de novo* purine biosynthesis. To summarize, at 8 hpi, the infected wound microenvironment has low purine metabolites **(Figure 2)** and high level of mannose **(Figure 4C)**. We found that purine availability is important to achieve high titers during the onset of *E. faecalis* wound infection. Therefore, our data suggests that *E. faecalis* lacking *mptD* would be attenuated due to its inability to overcome the low purine bioavailability by being unable to uptake mannose to trigger its own purine biosynthesis.

### *E. faecalis de novo* purine biosynthesis and MptABCD phosphotransferase system are important for catheter-associated urinary tract infection

Purine biosynthesis is essential in a variety of infection types (Goncheva et al., 2020; Kim et al., 2003; Li, L. et al., 2018; Mei et al., 1997; Samant et al., 2008; Sause et al., 2019), and we therefore wondered whether *E. faecalis de novo* purine biosynthesis and MptABCD PTS would similarly contribute to other *E. faecalis* infections. In addition to being a common wound pathogen, *E. faecalis* is also a frequently isolated uropathogen (Kline & Lewis, 2016). Hence, we tested the contribution of purine biosynthesis and MptABCD PTS in a CAUTI model, assessing *in vivo* competitive infections of OG1RF Δ*purEK* and OG1RF Δ*mptD* with OG1X at 1 dpi. We observed that OG1RF Δ*purEK* (CI = 8.99) and OG1RF Δ*mptD* (CI = 6.01) had significantly reduced fitness compared to wild-type OG1RF (CI = 46.19) on catheters **(Figure 6A)**, while only OG1RF Δ*mptD* (CI = 4.83) had significantly reduced fitness compared to wild-type OG1RF (CI = 33.33) in the bladder **(Figure 6B)**, and neither *purEK* nor *mptD* contributed to fitness in the kidneys **(Figure 6C)**. These results suggest that *de novo* purine biosynthesis and the MptABCD PTS may be central and niche-independent virulence factors of *E. faecalis*.

**Figure 6.**
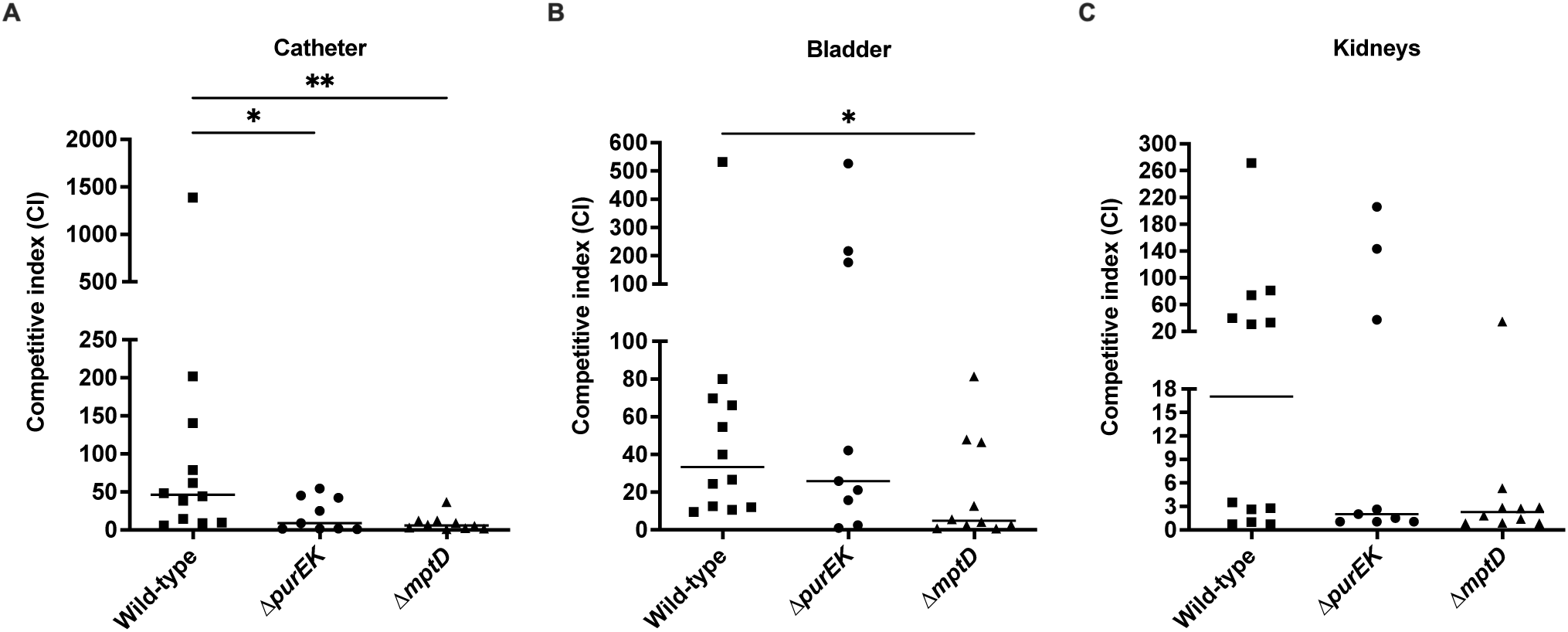
*De novo* purine biosynthesis and MptABCD phosphotransferase system contributes to *E. faecalis* fitness during CAUTI. Female C57BL/6 mice were implanted with 5 mm silicon catheters in the bladders and infected with a 1:1 ratio of 10^7^ CFU of *E. faecalis* OG1X:wild-type OG1RF, OG1X:OG1RF Δ*purEK* or OG1X:OG1RF Δ*mptD*. **(A)** Catheters, **(B)** bladders and **(C)** kidneys were harvested at 24 hpi and the recovered bacteria were enumerated on selective agar plates for each strain. Each data point represents one mouse and horizontal lines indicate the median; N = 2, n = 6 mice per group per experiment. Statistical analysis was performed using the Mann-Whitney U test, *p < 0.05, **p < 0.01.

## DISCUSSION

In this study, we sought to identify fitness determinants that are crucial for acute *E. faecalis* replication and later persistence in wounds. We show that both *E. faecalis de novo* purine biosynthesis and MptABCD PTS are important for *E. faecalis* acute replication and persistence, respectively. We report that purine metabolites are lower at the wound site due to rapid consumption by *E. faecalis* during the early stages of wound infection compared to later persistent stages, explaining the importance of *de novo* purine biosynthesis for acute *E. faecalis* wound infection. We also show that carbohydrate availability in the wound microenvironment has more galactose and mannose as the wound infection progresses, providing a reason for the requirement of the MptABCD galactose and mannose transporter during persistent *E. faecalis* wound infection.

Nucleotide plays a critical role in cell physiology of both prokaryotes and eukaryotes, such as DNA and RNA synthesis, enzyme cofactors (NAD^+^ and FAD^+^), energy carriers (ATP and GTP), and are also involved in the biosynthesis of riboflavin (Abbas & Sibirny, 2011; Jensen et al., 2008). *De novo* purine biosynthesis is required for many pathogens to establish a successful infection. For example, purine biosynthesis is necessary for successful proliferation of Gram-negative *E. coli* and *Salmonella typhimurium* in human serum (Samant et al., 2008) and *P. aeruginosa* in wounds (Turner et al., 2014) as well as for Gram-positive *Streptococcus pyogenes* growth in human blood (Le Breton et al., 2013), *Enterococcus faecium* growth in human serum (Zhang et al., 2017) and *Bacillus anthracis* growth in human serum and virulence in a mouse bacteremia model (Samant et al., 2008). Likewise, purine biosynthesis is required for *S. aureus* growth in bovine and human serum (Connolly et al., 2017), virulence in mouse models of bacteremia (Goncheva et al., 2020; Sause et al., 2019) and endocarditis infections (Li, L. et al., 2018). Therefore, it is not surprising that purine biosynthesis is also required for *E. faecalis* to establish a successful colonization in wounds, especially during the early phase of wound infection where *E. faecalis* is rapidly replicating (Chong et al., 2017). Together, these studies demonstrate that *de novo* purine biosynthesis probably is a common metabolic pathway that is generally required by bacterial pathogens in most infections. Hence, finding ways to locally sequester exogenous purines in the wound microenvironment might be useful in controlling *E. faecalis* colonization during wound infection.

Carbohydrates are essential for their metabolism into glucose, which serves as a primary energy source for most bacteria. These large uncharged polar molecules cannot cross the bacterial plasma membrane freely (Cooper & Hausman, 2000). Consequently, bacteria encode PTS to import carbohydrates from the environment (Deutscher et al., 2006). A PTS is made up of several functional subunits, of which the EII subunits of each PTS determines its substrate carbohydrate specificity (Deutscher et al., 2006). As such, most bacteria encode multiple PTS to enable the import of different carbohydrates. As the wound infection progresses, *E. faecalis* encounters changing carbohydrate availability in the wound microenvironment from higher glucose during acute wound infection to higher galactose and mannose in late stages of the infection, and because *E. faecalis* can persist and survive the change in carbohydrate availability, it hints that there is a metabolic switch in the carbohydrate phosphotransferase system in *E. faecalis* and that wound infection is a “controlled” process. As a result, any disturbance introduced (e.g. hindered mannose import) to this “controlled” process would then lead to reduced competitive index as observed with the OG1RF Δ*mptD* mutant during wound infection. However, there is still limited information on carbohydrate availability in wounds in other animal models or in human wounds, hence we cannot discount the presence of other carbohydrates in the wound microenvironment and the importance of other PTS that might be contributing to *E. faecalis* persistence. Apart from *E. faecalis*, the impact of carbohydrates metabolism and import is also evident in the pathogenesis of several other Gram-positive bacteria. For example, sucrose-6-phosphate hydrolase and its sucrose ABC transporter contributes to *Streptococcus pneumoniae in vivo* fitness during lung infection in mouse (Iyer & Camilli, 2007). Based on comparative genomic analysis, a PTS locus in *Enterococcus faecium* clinical isolates is found to play an important role in mouse intestinal colonization and the deletion of an EII subunit of this PTS resulted in reduced intestinal colonization (Zhang et al., 2013). Garnett et al. (2014) similarly showed the importance of *S. aureus* PTS in importing carbohydrates from the airway surface liquid to support its growth. Given the significance of PTS on the pathogenesis of various infections caused by different bacteria, drugs and/or inhibitors targeting carbohydrate import process(s) seems like an attractive alternative to control infections.

As aforementioned, purine biosynthesis may be a common metabolic pathway that is required for virulence. Fittingly, *E. faecalis de novo* purine biosynthesis also contributes to its fitness during CAUTI. The notion that purines are limiting in the urinary tract is consistent with studies of uropathogenic *E. coli*, in which a *guaA* mutant that has defective guanine biosynthesis was unable to grow in human urine *in vitro* and was significantly less virulent than the parental wild-type strain in a mouse model of urinary tract infection (UTI) (Russo et al., 1996). The OG1RF Δ*mptD* mutant was less fit than wild-type during CAUTI, suggesting that galactose and mannose availability in the urinary tract are likely limited, which is in contrast with uropathogenic *E. coli* that preferentially take advantage of amino acids and small peptides as a carbon source, since mutants with defective peptide import had significantly reduced fitness during UTI (Alteri et al., 2009). However, future studies will be needed to confirm whether purines are similarly limited, as well as the carbohydrate profile in the mice bladders during CAUTI.

*E. faecalis de novo* purine biosynthesis and MptABCD PTS are functionally linked **(Figure 5B)**. There are multiple pathways like the pentose phosphate pathway, alanine, aspartate and glutamate metabolism, thiamine metabolism, and histidine metabolism that contributes to purine biosynthesis (efi00230) (Kanehisa & Goto, 2000), and it is possible that in order to maintain healthy levels of purine to support active cell division during *E. faecalis* acute replication, the presence of all the contributing pathways are likely required. This could explain why there is a growth attenuation of OG1RF Δ*mptD* mutant in purine lacking medium as purine biosynthesis is affected in the absence of mannose transport. However, the OG1RF Δ*mptD* mutant is strongly outcompeted compared to OG1RF Δ*purEK* in wounds at 3 dpi, suggesting that the import of galactose and mannose has a more significant role as a carbon source for *E. faecalis* persistence in wounds other than for purine biosynthesis. These observations suggest that genes can have a functional shift depending on the state/phase during *E. faecalis* wound infection.

An outstanding question from this study is whether shikimate biosynthesis might be contributing to *E. faecalis* persistence in wounds. The production of secondary metabolites are usually not critical for cell growth, but instead serve as a survival strategy for organisms during adverse conditions likely triggered by the depletion of nutrients or environmental stress (Gokulan et al., 2014). The end product of the shikimate pathway is chorismate, which is essential for subsequent biosynthesis of aromatic amino acids such as phenylalanine, tryptophan and tyrosine as well as aromatic secondary metabolites (Averesch & Krömer, 2018; Herrmann & Weaver, 1999). For example, chorismate branching into the synthesis of para-aminobenzoic acid (PABA), which is a precursor for folate metabolism (Averesch & Krömer, 2018). Interestingly, Turner et al. (2014) not only showed that purines were unavailable to *P. aeruginosa* during wound infection, chorismate, phenylalanine, tyrosine, and PABA were also unavailable. Moreover, shikimate pathway intermediates are also potential substrates leading to other metabolic pathways (Herrmann & Weaver, 1999). Thereby, it is tempting to hypothesize that the shikimate pathway is important as it is a central metabolic route that leads to the production of other aromatic metabolites that might be essential for *E. faecalis* persistence in wounds. However, further studies will be needed to examine the role of *E. faecalis* shikimate biosynthesis during wound infection.

Based on our current findings, we propose a working model of *E. faecalis* wound infection dynamics **(Figure 7)**. During the early phase of wound infection, *E. faecalis* undergoes an acute replication and therefore, the demand for purines is high. However, purine metabolites in the wound microenvironment are rapidly consumed by *E. faecalis*, which makes the *E. faecalis de novo* purine biosynthesis indispensable for the acute replication **(Figure 7A)**. Carbohydrate availability during acute infection differs from persistence, in which glucose is higher while galactose and mannose are lower during earlier stages of infection. Despite these differences in availability, it is likely that *E. faecalis* can use all three carbohydrates to support its rapid growth. By contrast, during *E. faecalis* persistence in wounds, galactose and mannose availability are higher than during acute replication and that there was minimal depletion of glucose by *E. faecalis*, suggesting that galactose and mannose are a preferred carbohydrate source by *E. faecalis* during persistence **(Figure 7B)**.

**Figure 7.**
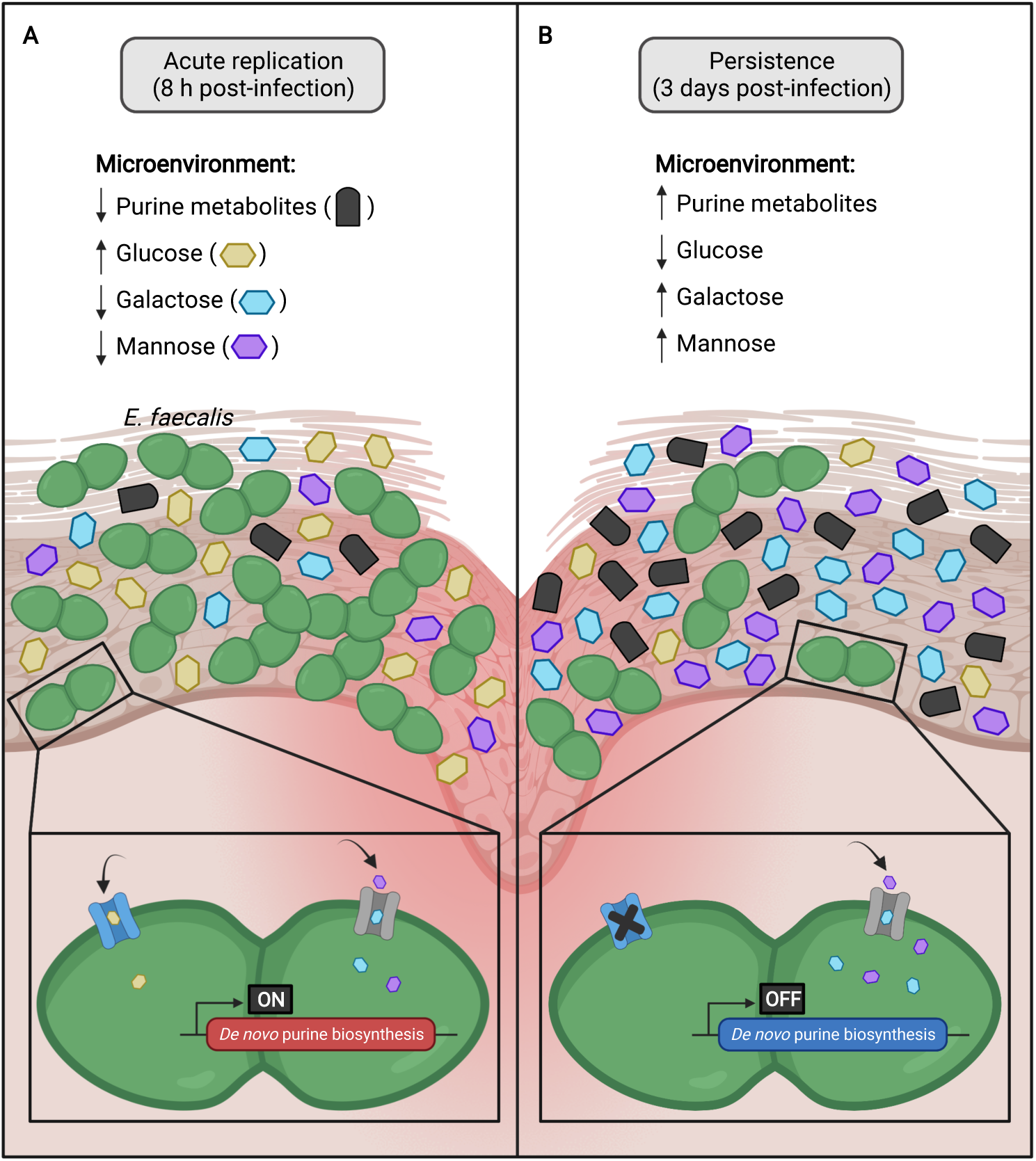
Proposed working model of *E. faecalis* wound infection dynamics. **(A)** During *E. faecalis* acute replication (8 h post-infection), purine metabolites, galactose and mannose availability in the wound microenvironment are low while glucose is high. As a result, *de novo* purine biosynthesis is induced and *E. faecalis* imports all three carbohydrates to support its rapid growth. **(B)** However, as the wound infection progresses to persistence (3 days post-infection), purine metabolites, galactose and mannose availability are high while glucose is low. Since *E. faecalis* is not actively dividing and purine metabolites are abundant in the wound microenvironment, *de novo* purine biosynthesis is likely not induced in *E. faecalis*. Additionally, given that there was minimal depletion of glucose and increased uptake of galactose and mannose by *E. faecalis*, it suggests that galactose and mannose are the preferred carbohydrates by *E. faecalis* during persistence. The figure was created with BioRender.com.

Overall, our study provides insights into the pathogenic requirements and potential of *E. faecalis* during wound infection, and factors that are required for *E. faecalis* to replicate and persist in this niche. Given the suggested importance of *E. faecalis de novo* purine biosynthesis and MptABCD PTS during acute replication and persistence in wounds, this work raises the possibility for future drugs and/or inhibitors to sequester exogenous purines in the wound microenvironment or to target MptABCD PTS, or in general *E. faecalis* carbohydrate utilization processes as a novel approach to curb infections.

## ACKOWLEDGEMENTS

This work was supported by the National Research Foundation and Ministry of Education Singapore under its Research Centre of Excellence Program and by a grant from the National Medical Research Council (MOH-OFIRG18may-005) awarded to K.A.K. This work was part supported by the Ministry of Education Singapore under its Singapore Ministry of Education Academic Research Fund Tier 1 (2019-T1-001-059) awarded to Y.A. Preparation of this article was also financially supported by the Interdisciplinary Graduate Program of Nanyang Technological University. We thank Gary M. Dunny from University of Minnesota Medical School for the *E. faecalis* transposon library.

---

**Supplementary Figure 1.**
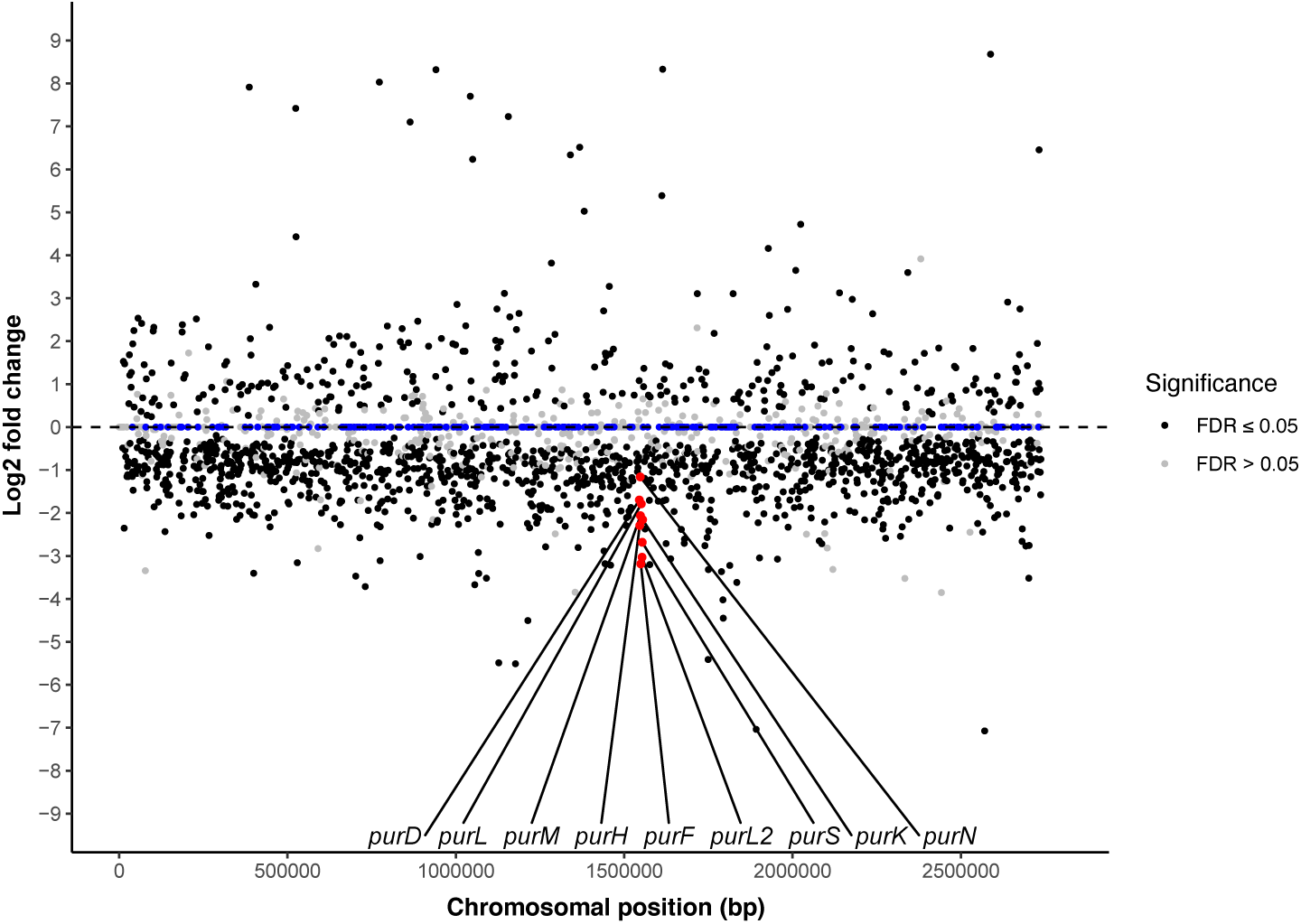
Transposon insertions in *E. faecalis de novo* purine biosynthesis genes are among the most significantly underrepresented genes at 8 hpi. Distribution of *E. faecalis* transposon mutant abundance profiled by Tn-seq from 8 hpi wounds. Significant mutants from Tn-seq analysis are colored black (p ≤ 0.05 and FDR ≤ 0.05). *E. faecalis* genes with no transposon mutant found in the transposon library are colored in blue.

**Supplementary Figure 2.**
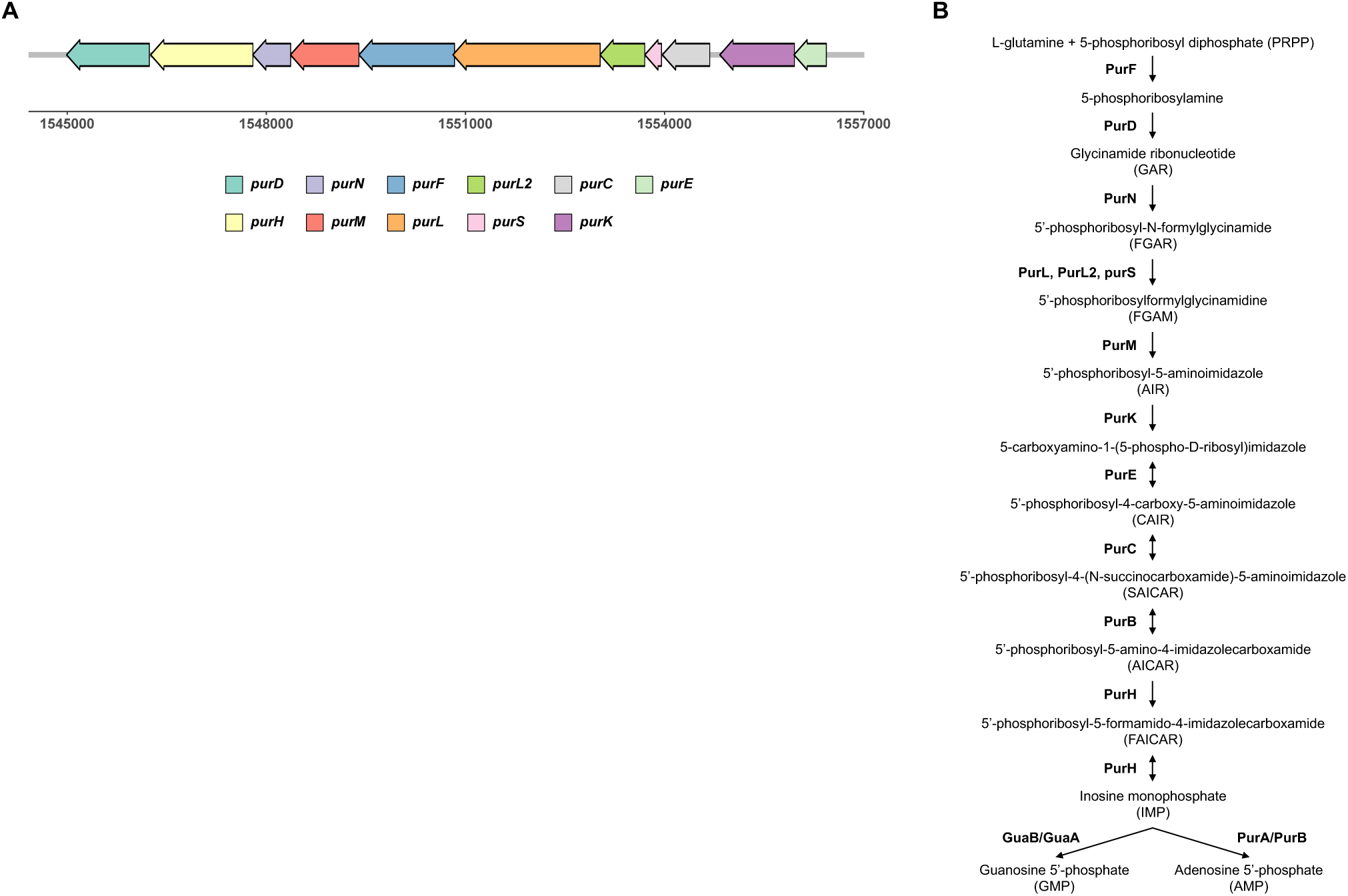
*De novo* purine biosynthesis in *E. faecalis.* **(A)** Operon arrangement of purine biosynthesis genes and **(B)** purine biosynthesis pathway in *E. faecalis*. Adapted from KEGG pathways efi00230.

**Supplementary Figure 3.**
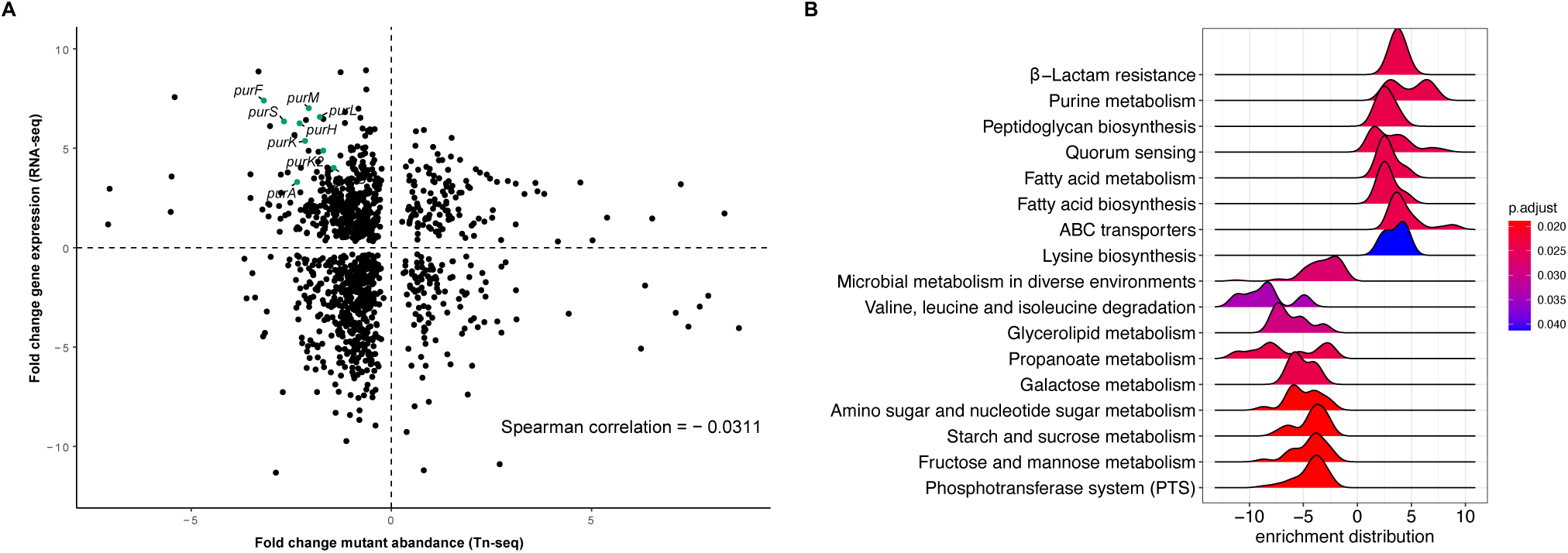
*E. faecalis* pathways that are significantly enriched in 8 hpi wounds. **(A)** Correlation plot of statistically significant genes from Tn-seq and RNA-seq. **(B)** Ridge plot showing the distribution of fold-change for genes in significantly enriched pathways identified from RNA-seq analysis. Color gradient represents false discovery rate (p.adjust values).

**Supplementary Figure 4.**
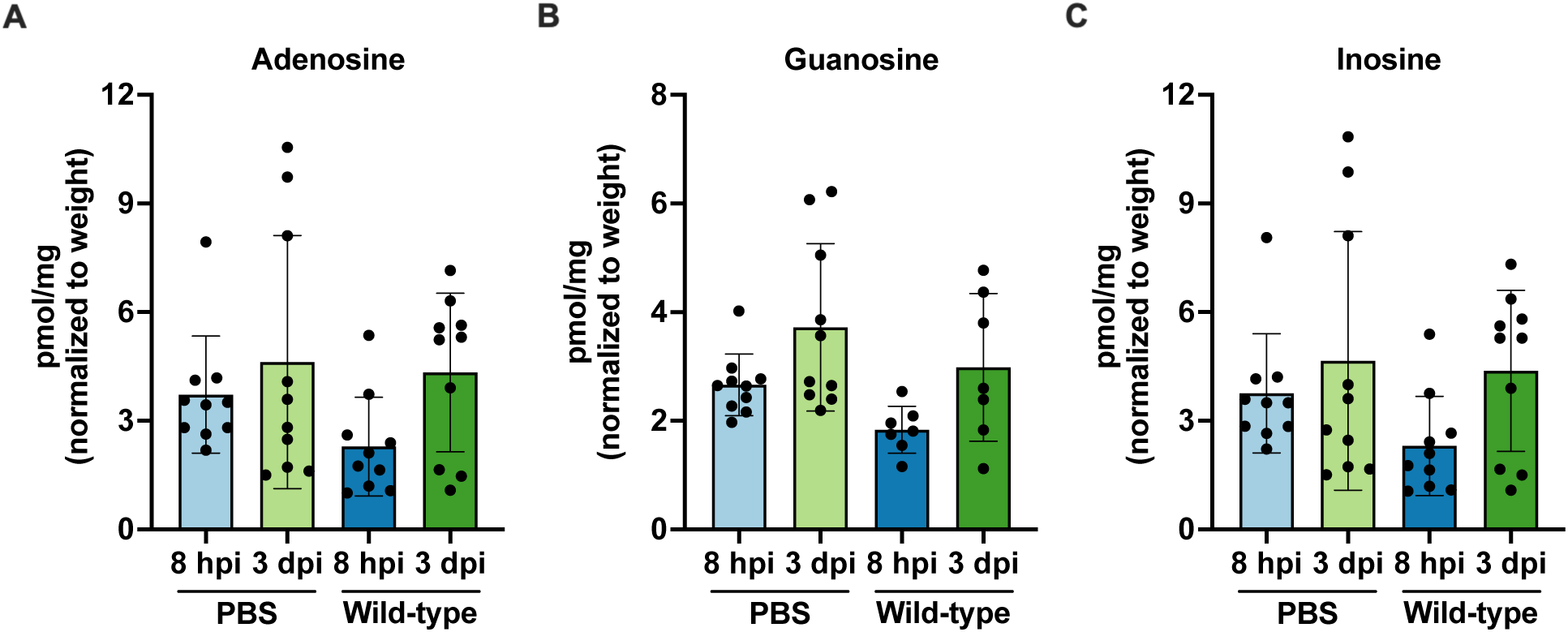
No significant differences between adenosine, guanosine, and inosine metabolite levels during *E. faecalis* wound infection. Male C57BL/6 mice were wounded and inoculated with PBS or 2 – 4 × 10^6^ CFU of wild-type OG1RF. Wounds were harvested at 8 hpi and 3 dpi for quantification of **(A)** adenosine, **(B)** guanosine, and **(C)** inosine using LC-MS. Each data point represents one mouse and error bars represent SD from the mean; N = 2, n = 5 mice per group per experiment. Statistical analysis was performed using the Mann-Whitney U test, *p < 0.05, **p < 0.01, ***p < 0.001.

**Supplementary Figure 5.**
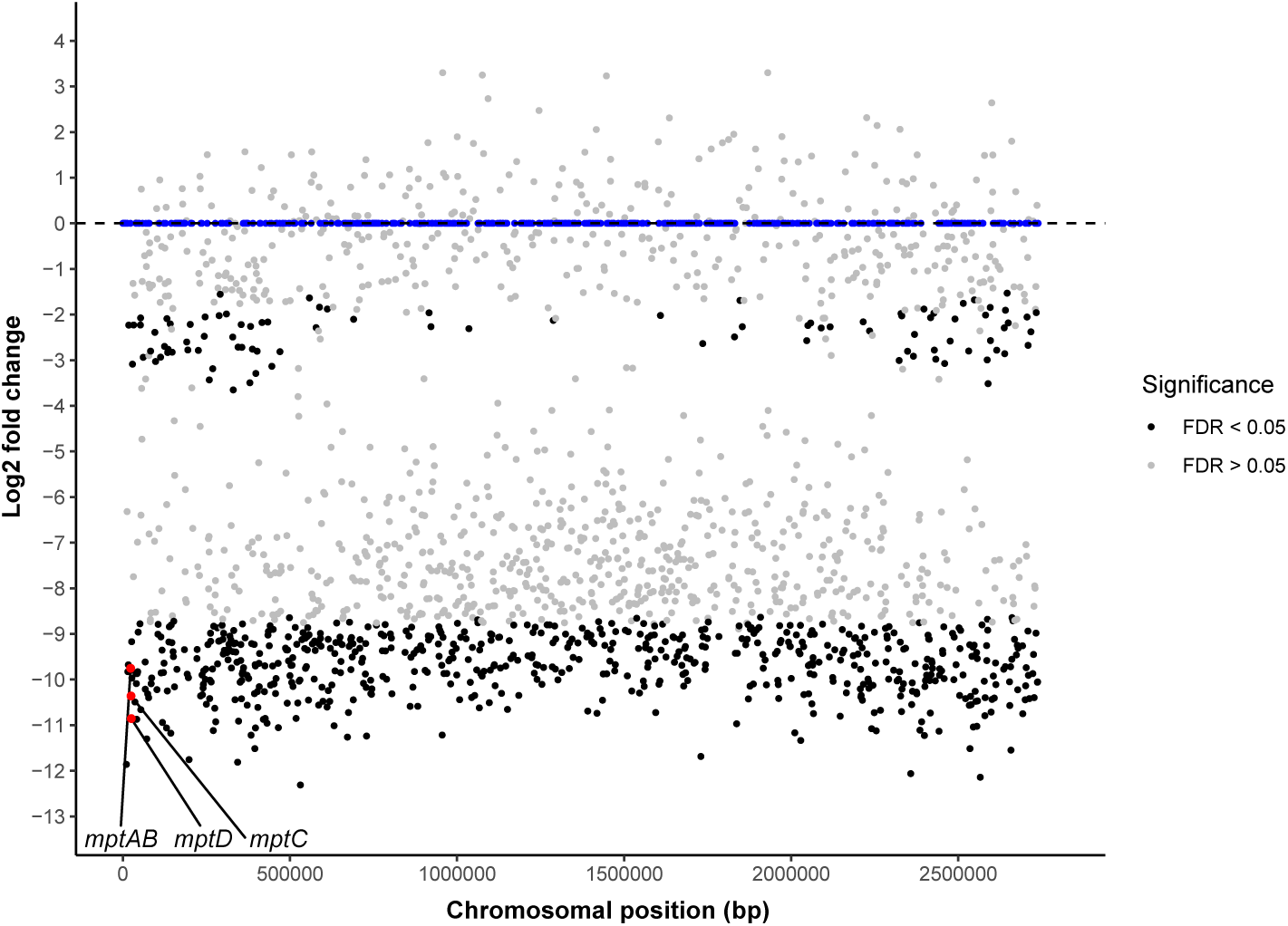
Transposon insertions in *mptABCD* are among the most significantly underrepresented genes at 3 dpi. Distribution of *E. faecalis* transposon mutant abundance profiled by Tn-seq from 3 dpi wounds. Significant mutants from Tn-seq analysis are colored black (p ≤ 0.05 and FDR ≤ 0.05). *E. faecalis* genes with no transposon mutant found in the transposon library are colored in blue.

**Supplementary Figure 6.**
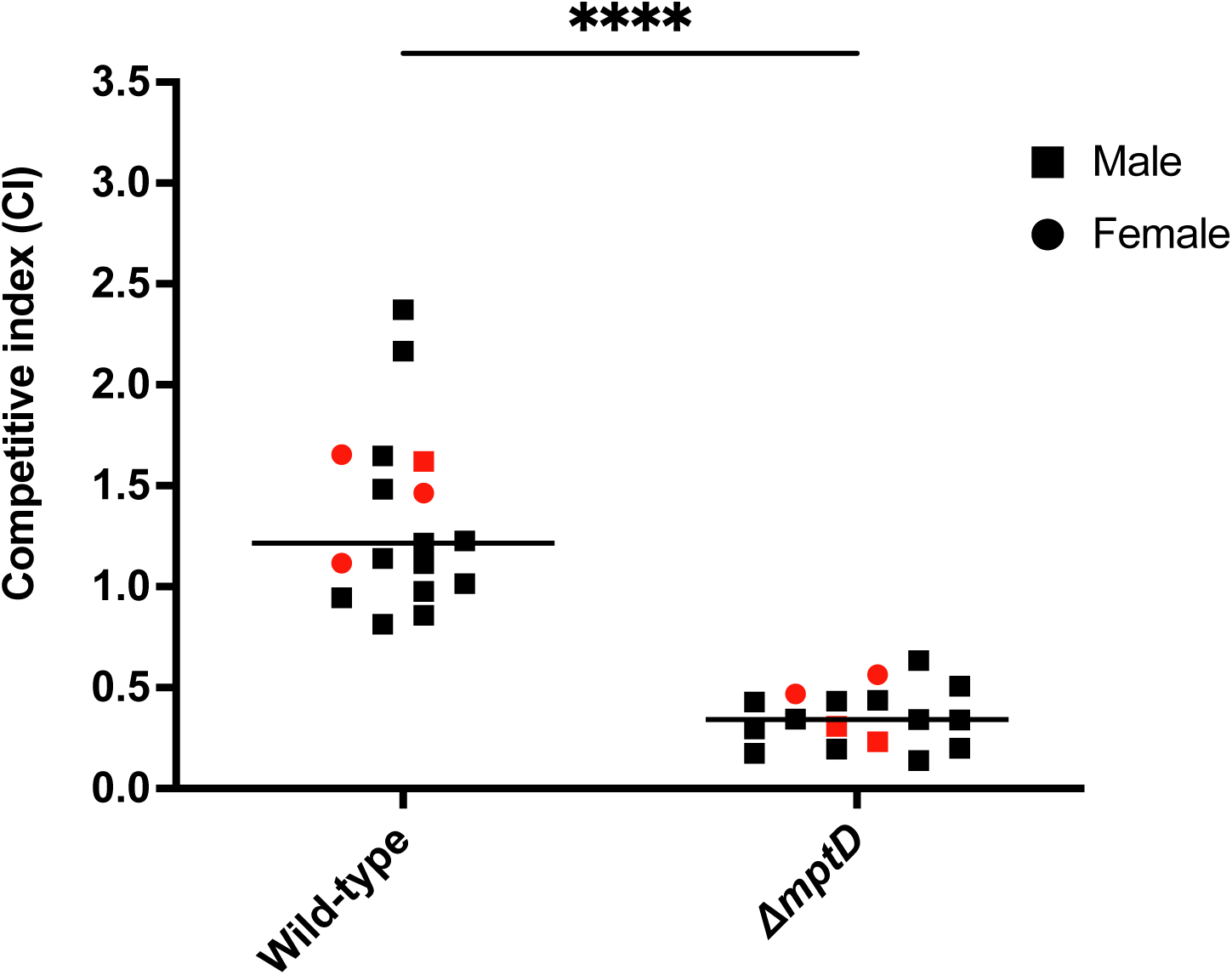
MptABCD phosphotransferase system contributes to *E. faecalis* wound fitness during persistence in diabetic mice. Male and female *db/db* mice were wounded and infected with a 1:1 ratio of *E. faecalis* OG1X:wild-type OG1RF or OG1X:OG1RF Δ*mptD* at 2 – 4 × 10^6^ CFU/wound (N = 4, n = 4 – 5 mice) and CFU determined at 3 dpi. The recovered bacteria were enumerated on selective agar plates for each strain. Each data point represents one mouse and horizontal lines indicate the median. Data points in red and black represents 7 – 8 weeks and 14 weeks old mice, respectively. Statistical analysis was performed using the Mann-Whitney U test, ****p < 0.0001.

**Supplementary Table 1.** Bacterial strains used in this study.

| Strain | Description | Reference |
| --- | --- | --- |
| OG1X | Wild-type, *str*<sup>r</sup> | Ike et al. (1983) |
| OG1RF | Wild-type, *rif*<sup>r</sup> | Dunny et al. (1978) |
| OG1RF pTCV::*P<sub>tet</sub>*-Empty | OG1RF with pTCV empty vector, *erm*<sup>r</sup>, *kan*<sup>r</sup> | This study |
| OG1RF pMSP3535::*P<sub>nisA</sub>*-Empty | OG1RF with pMSP3535 empty vector, *erm*<sup>r</sup> | This study |
| OG1RF Δ*purEK* | OG1RF with *purE* and *purK* deletion | This study |
| OG1RF Δ*purEK* pTCV::*P<sub>tet</sub>*-Empty | Δ*purEK* with pTCV empty vector, *erm*<sup>r</sup>, *kan*<sup>r</sup> | This study |
| OG1RF Δ*purEK* pTCV::*P<sub>tet</sub>*-*purEK* | Δ*purEK* with *purEK* complemented on pTCV vector, *erm*<sup>r</sup>, *kan*<sup>r</sup> | This study |
| OG1RF Δ*mptD* | OG1RF with *mptD* deletion | This study |
| OG1RF Δ*mptD* pMSP3535::*P<sub>nisA</sub>*-Empty | Δ*mptD* with pMSP3535 empty vector, *erm*<sup>r</sup> | This study |
| OG1RF Δ*mptD* pMSP3535::*P<sub>nisA</sub>*-*mptD* | Δ*mptD* with *mptD* complemented on pMSP3535 vector, *erm*<sup>r</sup> | This study |

*Str*<sup>r</sup>, *rif*<sup>r</sup>, *erm*<sup>r</sup> and *kan*<sup>r</sup> represents streptomycin, rifampicin, erythromycin, and kanamycin resistance, respectively.

**Supplementary Table 2.**
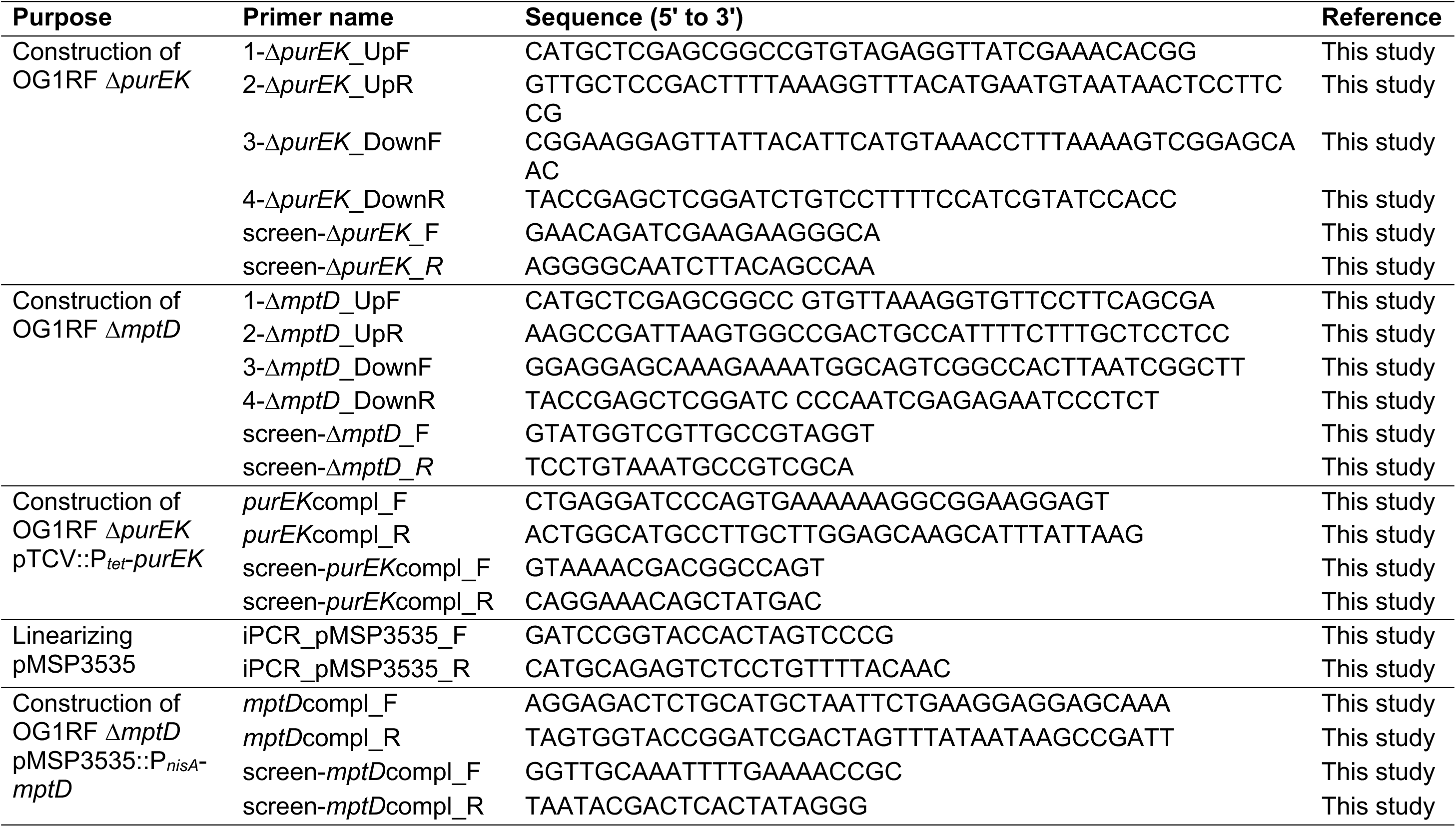
Primers used in this study.

**Supplementary Table 3.**
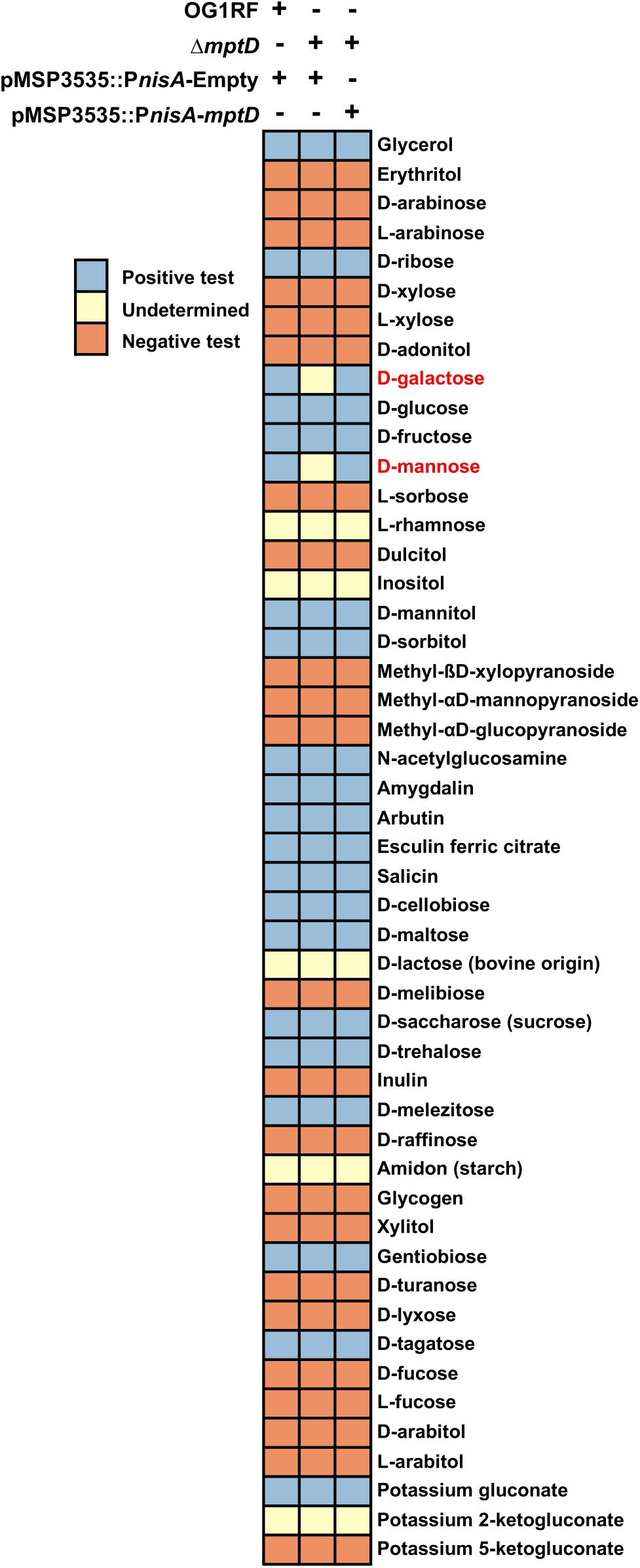
Complete table for carbohydrate fermentation test (API 50 CH) of wild-type OG1RF pMPSP3535::P*_nisA_*-Empty, OG1RF Δ*mptD* pMSP3535::P*_nisA_*-Empty, and OG1RF Δ*mptD* pMSP3535::*P_nisA_-mptD*.

## REFERENCES

Abbas, C. A., & Sibirny, A. A. (2011). Genetic control of biosynthesis and transport of riboflavin and flavin nucleotides and construction of robust biotechnological producers. Microbiology and Molecular Biology Reviews, 75(2), 321–360.

Alteri, C. J., Smith, S. N., & Mobley, H. L. (2009). Fitness of *Escherichia coli* during urinary tract infection requires gluconeogenesis and the TCA cycle. PLoS Pathogens, 5(5), e1000448.

Anders, S., Pyl, P. T., & Huber, W. (2015). HTSeq—a Python framework to work with high-throughput sequencing data. Bioinformatics, 31(2), 166–169.

Arias, C. A., & Murray, B. E. (2012). The rise of the *Enterococcus*: beyond vancomycin resistance. Nature Reviews Microbiology, 10(4), 266–278.

Averesch, N. J., & Krömer, J. O. (2018). Metabolic engineering of the shikimate pathway for production of aromatics and derived compounds—present and future strain construction strategies. Frontiers in Bioengineering and Biotechnology, 6, 32.

Bao, Y., Sakinc, T., Laverde, D., Wobser, D., Benachour, A., Theilacker, C., … Huebner, J. (2012). Role of *mprF1* and *mprF2* in the pathogenicity of *Enterococcus faecalis*. PloS One, 7(6), e38458.

Barquist, L., Mayho, M., Cummins, C., Cain, A. K., Boinett, C. J., Page, A. J., … Parkhill, J. (2016). The TraDIS toolkit: sequencing and analysis for dense transposon mutant libraries. Bioinformatics, 32(7), 1109–1111.

Bowler, P., Duerden, B., & Armstrong, D. G. (2001). Wound microbiology and associated approaches to wound management. Clinical Microbiology Reviews, 14(2), 244–269.

Bushnell, B. (2015). BBMap short-read aligner, and other bioinformatics tools. University of California, Berkeley, CA.

Chong, K. K. L., Tay, W. H., Janela, B., Yong, A. M. H., Liew, T. H., Madden, L., … Becker, D. L. (2017). *Enterococcus faecalis* modulates immune activation and slows healing during wound infection. The Journal of Infectious Diseases, 216(12), 1644–1654.

Connolly, J., Boldock, E., Prince, L. R., Renshaw, S. A., Whyte, M. K., & Foster, S. J. (2017). Identification of *Staphylococcus aureus* factors required for pathogenicity and growth in human blood. Infection and Immunity, 85(11), e00337–00317.

Cooper, G. M., & Hausman, R. (2000). A molecular approach. The Cell. *2nd ed*. *Sunderland, MA*: *Sinauer Associates*.

Dale, J. L., Beckman, K. B., Willett, J. L., Nilson, J. L., Palani, N. P., Baller, J. A., … Manias, D. A. (2018). Comprehensive functional analysis of the *Enterococcus faecalis* core genome using an ordered, sequence-defined collection of insertional mutations in strain OG1RF. MSystems, 3(5), e00062–00018.

Demidova-Rice, T. N., Hamblin, M. R., & Herman, I. M. (2012). Acute and impaired wound healing: pathophysiology and current methods for drug delivery, part 1: normal and chronic wounds: biology, causes, and approaches to care. Advances in Skin & Wound Care, 25(7), 304.

Deutscher, J., Francke, C., & Postma, P. W. (2006). How phosphotransferase system-related protein phosphorylation regulates carbohydrate metabolism in bacteria. Microbiology and Molecular Biology Reviews, 70(4), 939–1031.

Dowd, S. E., Sun, Y., Secor, P. R., Rhoads, D. D., Wolcott, B. M., James, G. A., & Wolcott, R. D. (2008). Survey of bacterial diversity in chronic wounds using Pyrosequencing, DGGE, and full ribosome shotgun sequencing. BMC Microbiology, 8(1), 43.

Dunny, G. M., Brown, B. L., & Clewell, D. B. (1978). Induced cell aggregation and mating in *Streptococcus faecalis*: evidence for a bacterial sex pheromone. Proceedings of the National Academy of Sciences, 75(7), 3479–3483.

Dworniczek, E., Piwowarczyk, J., Bania, J., Kowalska-Krochmal, B., Wałecka, E., Seniuk, A., … Gościniak, G. (2012). *Enterococcus* in wound infections: virulence and antimicrobial resistance. Acta Microbiologica et Immunologica Hungarica, 59(2), 263–269.

Fisher, K., & Phillips, C. (2009). The ecology, epidemiology and virulence of *Enterococcus*. Microbiology, 155(6), 1749–1757.

Flahaut, S., Hartke, A., Giard, J.-C., Benachour, A., Boutibonnes, P., & Auffray, Y. (1996). Relationship between stress response towards bile salts, acid and heat treatment in *Enterococcus faecalis*. FEMS Microbiology Letters, 138(1), 49–54.

Garnett, J. P., Braun, D., McCarthy, A. J., Farrant, M. R., Baker, E. H., Lindsay, J. A., & Baines, D. L. (2014). Fructose transport-deficient *Staphylococcus aureus* reveals important role of epithelial glucose transporters in limiting sugar-driven bacterial growth in airway surface liquid. Cellular and Molecular Life Sciences, 71(23), 4665–4673.

Giacometti, A., Cirioni, O., Schimizzi, A., Del Prete, M., Barchiesi, F., D’errico, M., … Scalise, G. (2000). Epidemiology and microbiology of surgical wound infections. Journal of Clinical Microbiology, 38(2), 918–922.

Gokulan, K., Khare, S., & Cerniglia, C. (2014). Metabolic pathways | Production of secondary metabolites of bacteria. In C. A. Batt & M. L. Tortorello (Eds.), Encyclopedia of Food Microbiology (Second Edition) (pp. 561–569). Oxford: Academic Press.

Goncheva, M. I., Flannagan, R. S., & Heinrichs, D. E. (2020). *De novo* purine biosynthesis is required for intracellular growth of *Staphylococcus aureus* and for the hypervirulence phenotype of a *purR* mutant. Infection and Immunity, 88(5), e00104–00120.

Guiton, P. S., Hung, C. S., Hancock, L. E., Caparon, M. G., & Hultgren, S. J. (2010). Enterococcal biofilm formation and virulence in an optimized murine model of foreign body-associated urinary tract infections. Infection and Immunity, 78(10), 4166–4175.

Herrmann, K. M., & Weaver, L. M. (1999). The shikimate pathway. Annual Review of Plant Biology, 50(1), 473–503.

Ike, Y., Craig, R. A., White, B. A., Yagi, Y., & Clewell, D. B. (1983). Modification of *Streptococcus faecalis* sex pheromones after acquisition of plasmid DNA. Proceedings of the National Academy of Sciences, 80(17), 5369–5373. doi:10.1073/pnas.80.17.5369

Iyer, R., & Camilli, A. (2007). Sucrose metabolism contributes to *in vivo* fitness of *Streptococcus pneumoniae*. Molecular Microbiology, 66(1), 1–13.

James, G. A., Swogger, E., Wolcott, R., Pulcini, E. d., Secor, P., Sestrich, J., … Stewart, P. S. (2008). Biofilms in chronic wounds. Wound Repair and Regeneration, 16(1), 37–44.

Jensen, K. F., Dandanell, G., Hove-Jensen, B., & WillemoËs, M. (2008). Nucleotides, nucleosides, and nucleobases. EcoSal Plus, 3(1).

Kandaswamy, K., Liew, T. H., Wang, C. Y., Huston-Warren, E., Meyer-Hoffert, U., Hultenby, K., … Henriques-Normark, B. (2013). Focal targeting by human β-defensin 2 disrupts localized virulence factor assembly sites in *Enterococcus faecalis*. Proceedings of the National Academy of Sciences, 110(50), 20230–20235.

Kanehisa, M., & Goto, S. (2000). KEGG: kyoto encyclopedia of genes and genomes. Nucleic Acids Research, 28(1), 27–30.

Kilstrup, M., Hammer, K., Ruhdal Jensen, P., & Martinussen, J. (2005). Nucleotide metabolism and its control in lactic acid bacteria. FEMS Microbiology Reviews, 29(3), 555–590.

Kim, Y. R., Lee, S. E., Kim, C. M., Kim, S. Y., Shin, E. K., Shin, D. H., … Hillman, J. D. (2003). Characterization and pathogenic significance of *Vibrio vulnificus* antigens preferentially expressed in septicemic patients. Infection and Immunity, 71(10), 5461–5471.

Kline, K. A., & Lewis, A. L. (2016). Gram-positive uropathogens, polymicrobial urinary tract infection, and the emerging microbiota of the urinary tract. Microbiology Spectrum, 4(2), 4.2. 04.

Kristich, C. J., Nguyen, V. T., Le, T., Barnes, A. M., Grindle, S., & Dunny, G. M. (2008). Development and use of an efficient system for random mariner transposon mutagenesis to identify novel genetic determinants of biofilm formation in the core *Enterococcus faecalis* genome. Applied and Environmental Microbiology, 74(11), 3377–3386.

Le Breton, Y., Mistry, P., Valdes, K. M., Quigley, J., Kumar, N., Tettelin, H., & McIver, K. S. (2013). Genome-wide identification of genes required for fitness of group A *Streptococcus* in human blood. Infection and Immunity, 81(3), 862–875.

Li, H., & Durbin, R. (2009). Fast and accurate short read alignment with Burrows– Wheeler transform. Bioinformatics, 25(14), 1754–1760.

Li, L., Abdelhady, W., Donegan, N. P., Seidl, K., Cheung, A., Zhou, Y.-F., … Xiong, Y. Q. (2018). Role of purine biosynthesis in persistent methicillin-resistant *Staphylococcus aureus* infection. The Journal of Infectious Diseases, 218(9), 1367–1377.

Mei, J. M., Nourbakhsh, F., Ford, C. W., & Holden, D. W. (1997). Identification of *Staphylococcus aureus* virulence genes in a murine model of bacteraemia using signature-tagged mutagenesis. Molecular Microbiology, 26(2), 399–407.

Nielsen, H. V., Guiton, P. S., Kline, K. A., Port, G. C., Pinkner, J. S., Neiers, F., … Hultgren, S. J. (2012). The metal ion-dependent adhesion site motif of the *Enterococcus faecalis* EbpA pilin mediates pilus function in catheter-associated urinary tract infection. mBio, 3(4), e00177–00112.

Peschel, A., Jack, R. W., Otto, M., Collins, L. V., Staubitz, P., Nicholson, G., … Tarkowski, A. (2001). *Staphylococcus aureus* resistance to human defensins and evasion of neutrophil killing via the novel virulence factor MprF is based on modification of membrane lipids with l-lysine. The Journal of Experimental Medicine, 193(9), 1067–1076.

Ramsey, M., Hartke, A., & Huycke, M. (2014). The physiology and metabolism of enterococci. Enterococci: From Commensals to Leading Causes of Drug Resistant Infection [Internet]: Massachusetts Eye and Ear Infirmary.

Robinson, M. D., McCarthy, D. J., & Smyth, G. K. (2010). edgeR: a Bioconductor package for differential expression analysis of digital gene expression data. Bioinformatics, 26(1), 139–140.

Russo, T. A., Jodush, S. T., Brown, J. J., & Johnson, J. R. (1996). Identification of two previously unrecognized genes (*guaA* and *argC*) important for uropathogenesis. Molecular Microbiology, 22(2), 217–229.

Samant, S., Hsu, F.-F., Neyfakh, A. A., & Lee, H. (2009). The *Bacillus anthracis* protein MprF is required for synthesis of lysylphosphatidylglyceros and for resistance to cationic antimicrobial peptides. Journal of Bacteriology, 191(4), 1311–1319.

Samant, S., Lee, H., Ghassemi, M., Chen, J., Cook, J. L., Mankin, A. S., & Neyfakh, A. A. (2008). Nucleotide biosynthesis is critical for growth of bacteria in human blood. PLoS Pathogens, 4(2), e37.

Sause, W. E., Balasubramanian, D., Irnov, I., Copin, R., Sullivan, M. J., Sommerfield, A., … Ueberheide, B. (2019). The purine biosynthesis regulator PurR moonlights as a virulence regulator in *Staphylococcus aureus*. Proceedings of the National Academy of Sciences, 116(27), 13563–13572.

Sen, C. K., Gordillo, G. M., Roy, S., Kirsner, R., Lambert, L., Hunt, T. K., … Longaker, M. T. (2009). Human skin wounds: a major and snowballing threat to public health and the economy. Wound Repair and Regeneration, 17(6), 763–771.

Shettigar, K., Bhat, D. V., Satyamoorthy, K., & Murali, T. S. (2018). Severity of drug resistance and co-existence of *Enterococcus faecalis* in diabetic foot ulcer infections. Folia Microbiologica, 63(1), 115–122.

Thedieck, K., Hain, T., Mohamed, W., Tindall, B. J., Nimtz, M., Chakraborty, T., … Jänsch, L. (2006). The MprF protein is required for lysinylation of phospholipids in listerial membranes and confers resistance to cationic antimicrobial peptides (CAMPs) on *Listeria monocytogenes*. Molecular Microbiology, 62(5), 1325–1339.

Turner, K. H., Everett, J., Trivedi, U., Rumbaugh, K. P., & Whiteley, M. (2014). Requirements for *Pseudomonas aeruginosa* acute burn and chronic surgical wound infection. PLoS Genetics, 10(7).

Wu, T., Hu, E., Xu, S., Chen, M., Guo, P., Dai, Z., … Yu, G. (2021). clusterProfiler 4.0: A universal enrichment tool for interpreting omics data. Innovation (N Y), 2(3), 100141. doi:10.1016/j.xinn.2021.100141

Yu, G., Wang, L. G., Han, Y., & He, Q. Y. (2012). clusterProfiler: an R package for comparing biological themes among gene clusters. OMICS, 16(5), 284–287. doi:10.1089/omi.2011.0118

Zhang, X., de Maat, V., Guzmán Prieto, A. M., Prajsnar, T. K., Bayjanov, J. R., de Been, M., … Willems, R. J. (2017). RNA-seq and Tn-seq reveal fitness determinants of vancomycin-resistant *Enterococcus faecium* during growth in human serum. BMC Genomics, 18(1), 1–12.

Zhang, X., Top, J., de Been, M., Bierschenk, D., Rogers, M., Leendertse, M., … van Schaik, W. (2013). Identification of a genetic determinant in clinical *Enterococcus faecium* strains that contributes to intestinal colonization during antibiotic treatment. The Journal of Infectious Diseases, 207(11), 1780–1786.

